# Neural representation of action grammar structure in primate frontal cortex

**DOI:** 10.64898/2026.01.18.700235

**Authors:** Lucas Y. Tian, Daniel J. Hanuska, Kedar Garzón Gupta, Yue Liu, Xiao-Jing Wang, Joshua B. Tenenbaum, Winrich A. Freiwald

## Abstract

Humans and other animals rapidly solve novel problems, often on the first attempt. This capacity is hypothesized to rely on grammars, or internalized systems of abstract rules for recombining units of knowledge into novel composites. Yet, whether and how the brain realizes grammars remains unclear. Here we identify a neural population that represents the latent structure of a grammar. Macaque monkeys traced complex geometric figures with stroke sequences according to the action grammar A^n^B^m^C^k^, meaning “repeat A, then B, then C”, where A, B and C are variables that take specific shape values (e.g., A=circle), and n, m, and k correspond to numbers of repetitions. Macaques used this learned grammar to solve unfamiliar problems, including ones more complex than those seen in training—evidence for immediate grammar-based generalization. Recordings across eight frontal cortical areas revealed a representation of grammar structure during drawing, strongest in the presupplementary motor area (preSMA). A, B, and C were represented as distinct states that abstract over specific shapes, resembling roles in hypothesized role-filler structures underlying symbolic cognition. Ordinal position during repetition was encoded along linear axes that were largely orthogonal across A, B, and C. Thus, preSMA represents the geometry of action grammar structure supporting the use of internalized rules to generate novel action sequences.

## Introduction

Humans and other animals can solve problems by constructing solutions that recombine known elements in novel ways^1–5^. This capacity for compositional generalization^2,6–9^ is hypothesized to depend on grammars^2,4,7,10–14^—internalized rule systems that can be applied to combine units of knowledge into composites. Because a finite set of rules in a grammar can in principle be composed (e.g., by applying rules in sequence) to generate a large, potentially unbounded, set of possible solutions, grammars provide a potential explanation for immediate (or “zero-shot”) generalization to solving novel problems. Beyond the obvious case of language understanding, grammars have been hypothesized for a wide range of abilities^4,11,12,15–26^, from mathematical and logical reasoning^11,15,26–28^ to less overtly structured abilities such as tool use and object manipulation^17,25^, musical ability^11^, handwriting^18^, drawing^16,29^, foraging^30^, communication^20,24,31^, and action sequencing^4,10,32,33^. For example, when asked to “draw an animal that doesn’t exist,” children readily produce imaginary animals, such as a dog-like creature with six legs and three camel humps^19^, consistent with knowledge of rules like “repeat body part X” or “attach part X to part Y.”

Despite this behavioral evidence, whether and how grammars are realized in the brain remains unclear. A fundamental question is whether neural activity explicitly represents the abstract structure of grammars independent from specific content, such as specific objects, events, or actions^7,11,34–37^. For example, activity may reflect the abstraction “repeat body part X and then attach to Y” the same way whether X is a head or an arm. Representing abstract structure is useful because it allows structure-level operations to apply across arbitrary content, thereby supporting generalization across different situations involving the same rules. Accordingly, the strong capacities for generalization in language and other cognitive behaviors^2,11,13,38^ have led to the hypothesis that the brain represents abstract grammar structures^7,34,39,40^. This idea has been formalized in neural network models of cognition^36,41,42^ and is supported by evidence from human functional imaging studies^7^. However, evidence in neural activity for representation of grammar structure—abstracted from content and used during grammar-based problem solving—has been limited. Critically, it is not a given that such structure must exist in neural activity. For instance, modern artificial intelligence (AI) systems exhibit advanced language and cognitive abilities despite being built on distributed processing in neural network architectures rather than on explicit implementation of grammatical rules. This raises the possibility that the brain may similarly rely on mechanisms lacking explicit representation of grammar structure^43,44^.

A key challenge in studying grammars in neural activity has been the lack of an experimental paradigm that combines grammar-based problem-solving with neural recordings. To establish such a paradigm, we developed an action sequencing task satisfying three properties we believe to be important for studying grammars. First, behavior should be internally generated, in that the subject must decide, with no instructive cues, how to solve any given problem by applying a general set of rules. Second, behavior should generalize, including to harder, novel problems the first time they are encountered. Third, the task should allow experimentally dissociating structure from content to test for abstraction. This involves having different problems that use the same rules (e.g., repeat body part X) with different content (e.g., X = head or X = arm).

To our knowledge, prior studies combining neuronal recordings and behavior have not combined these three properties. However, multiple lines of study have yielded relevant insights. First, studies of “artificial grammar learning”^4,45,46^ have found that many species can discriminate grammatically structured regularities in stimulus sequences (e.g., sequences of sounds); however, these studies have typically relied on tasks requiring perceptual decisions rather than generation of grammar-based behavior. Second, recordings during action sequencing have revealed diverse neural correlates, but these have been in tasks involving either explicit ordering cues^47–52^, memorized sequences^53–57^, or rules but without assessing the ability to apply them on novel problems^50,52,58–62^. Third, some studies have involved rules applied to novel problems, but either have not demonstrated immediate generalization^49^ or have lacked neural recordings^32,33,63–66^. Fourth, ethological studies have described grammars in behaviors such as object manipulation^17^, foraging^30^, and tool use^67^, but these settings raise difficulties for experimental control, testing for novel behavior, and recording neural activity. In sum, although abstract properties have been found in neural activity for a variety of sequencing paradigms^49,50,52,54–59,61,68–71^, we currently lack a task that probes abstract grammar structure in behavior with simultaneous neural recordings.

We addressed this gap by training macaque monkeys to perform a drawing-like task involving grammar-based action sequencing, combined with neural recordings across eight frontal cortical areas previously implicated in rule use or action sequencing, but never directly compared. We found a dominant role for the presupplementary motor area (preSMA) in supporting this grammatical ability. During drawing, preSMA population activity encodes the abstract relationship between the ongoing action and the overall latent structure of the A^n^B^m^C^k^ grammar.

## Results

### Curriculum-based training of grammar-based visual-motor construction

We trained monkeys on a drawing-like task requiring them to reconstruct images of complex geometric figures by tracing with a sequence of strokes. Sequencing had to follow a hierarchical repetition grammar having the structure A^n^B^m^C^k^, meaning “repeat A, then repeat B, then repeat C”, where A, B, and C are roles, or abstract variables that can be assigned arbitrary shape values through learning (e.g., A=circle), and where the number of repetitions of each role (n, m, and k) vary across images (general grammar structure in **Fig. 1a**; example experiment in **Fig. 1b, c**). One stroke was allowed per instance of each role (**Fig. 1d, e**; touchscreen setup in **Extended Data Fig. 1**), and each role’s shape was drawn using its associated stroke type (action symbol) learned in a previous study^5^ (**Fig. 1a**, “Shapes”). This grammar was inspired by grammatical structure seen in human drawing^16^. In general, each role was associated with only one shape for each experimental session. In a specific case, each role was associated with two shapes, switched between on a trial-by-trial basis, to test for abstraction of role across specific shapes (**Fig. 4**).

**Figure 1.**
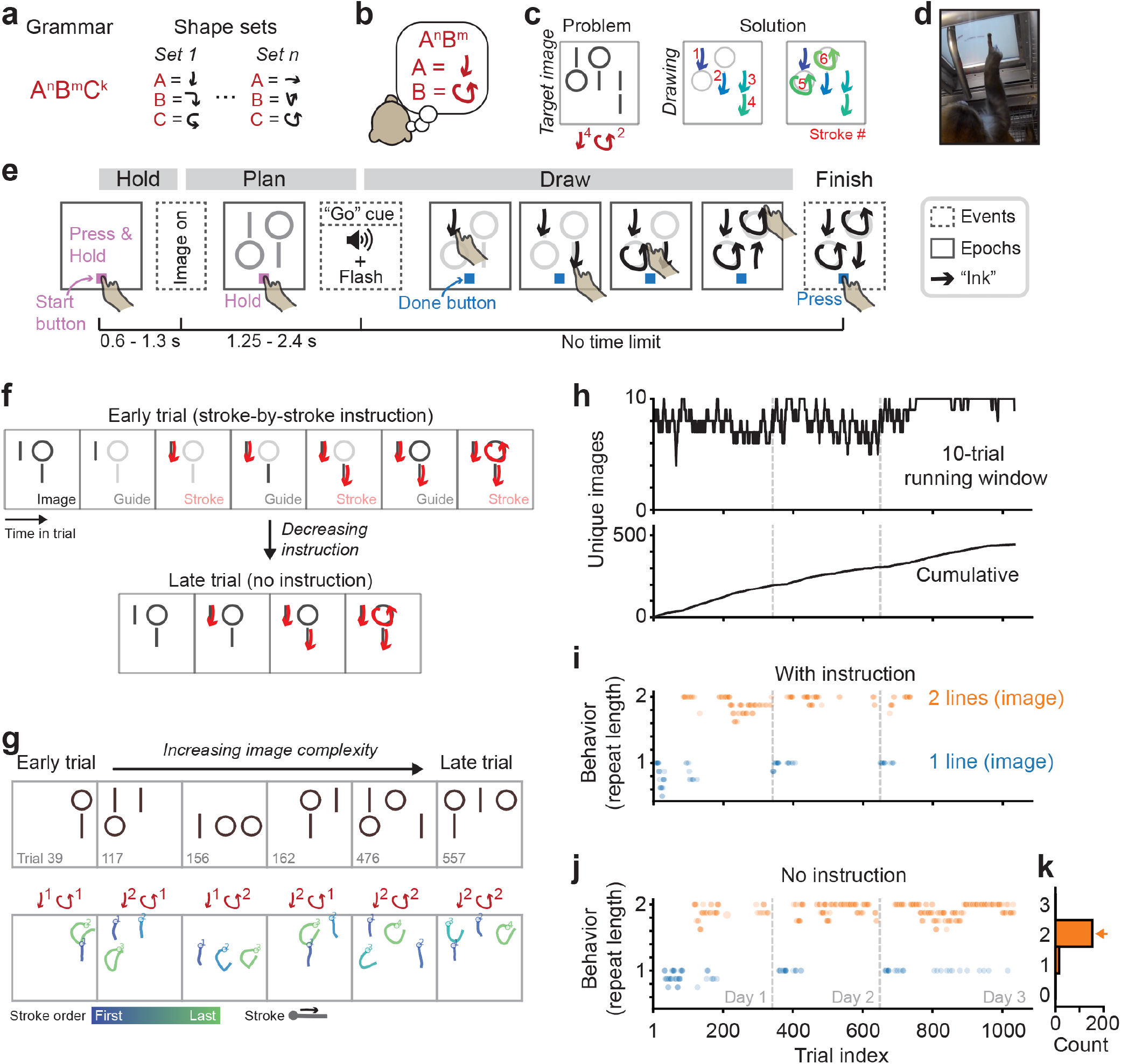
Curriculum-based training of grammar-based visual-motor construction. (a) The general form of action grammars in our study (top-left). The roles A, B, and C were each associated with a specific shape through training (“Shape sets”). The current study involved two variants of the grammar task, one in which one set of role-shape associations (i.e., shape set) was learned per experiment, and another in which two shape sets were switched between on a trial-by-trial basis, cued by the shape content of the image (Fig. 4). (b) The specific action grammar for the experiment in panels b-k. Within each role repeat, the correct direction to order strokes is from left to right. (c) Schematic of the core idea of the task. The subject’s goal is to apply a learned grammar to determine the correct stroke sequence for any problem (image). For this example problem, the correct sequence is to repeat “line” four times and then repeat “circle” twice, drawing left-to-right within each shape’s repeat. (d) Photo during in-cage training (the metal tube is the reward spout). (e) Trial structure, showing three discrete events (dashed boxes) and sustained epochs (solid boxes). “Hold”: press and hold a “start” button, enforcing a consistent posture. “Plan”: subjects see the image but must maintain hold on “start” button. “Draw”: subjects produce strokes. Black arrows represent “ink” (visible to the subject) marking where strokes have been produced. Subjects can report completion any time by pressing “done” button. They receive juice reward from producing a sequence consistent with the trained grammar (Methods). “Buttons” always refers to touchscreen buttons. (f) Schematic of one component of the training curriculum: a decrease in stroke-by-stroke instruction over trials. Instruction involved cuing the correct stroke one-by-one. Instruction was decreased through stages in a performance dependent manner by gradually lowering the difference in contrast between the next item and other items, until this difference reached zero, thus removing instruction. (g) Schematic of the second component of the training curriculum, an increase in image complexity over trials, which occurred in parallel with the decrease in instruction (panel f). Image complexity was defined as the number of total shape components, which, in this experiment, was increased from two to four (maximum two circles, two lines), in a manner depending on performance. (h) The number of unique images (including those that vary only by translation) counted in either a 10-trial running window (top) or cumulatively over all trials (bottom). (i) Behavior over three days of training, for trials with stroke-by-stroke instruction present, quantified as the running average (bin size of 8 trials) of the number of repeated lines (role *A*), plotted separately for images with one (blue) and two (orange) lines. For this experiment’s A^n^B^m^ grammar, we focused analysis on role A, because this is a stronger test of grammar understanding than analysis of role B, because the correct repeat length for role B (and in general, the last role) can be attained by simply producing the image components that have not been drawn, a strategy which does not require the grammar (Methods). (j) Same as panel i, but for trials without instruction. (k) Summary of behavior (repeat length) without instruction for two-line images on the last day of training (day 3). Orange arrow, expected length for correct performance.

Curriculum training is detailed for an example experiment (**Fig. 1b-k**) in which the monkey was trained on the grammar A^n^B^m^ with one shape set (A=line, B=circle; **Fig. 1b**). On each trial, the monkey was presented with a problem in the form of a static image composed of 0-5 lines and 0-5 circles. For training problems (0-2 lines and 0-2 circles) reward was provided at the end of the trial for producing a sequence that was correct according to the grammar. In general, we ensured each problem had only one correct sequence by imposing a direction rule (e.g., left-to-right in **Fig. 1b-k**; in a few experiments we used the subject’s default direction rather than imposing a rule; Methods). Test problems were designed to be harder by having more shape components (3-5 lines, 3-5 circles; **Fig. 2**). A correct sequence for an example test problem with four lines and two circles is shown in **Fig. 1c**.

**Figure 2.**
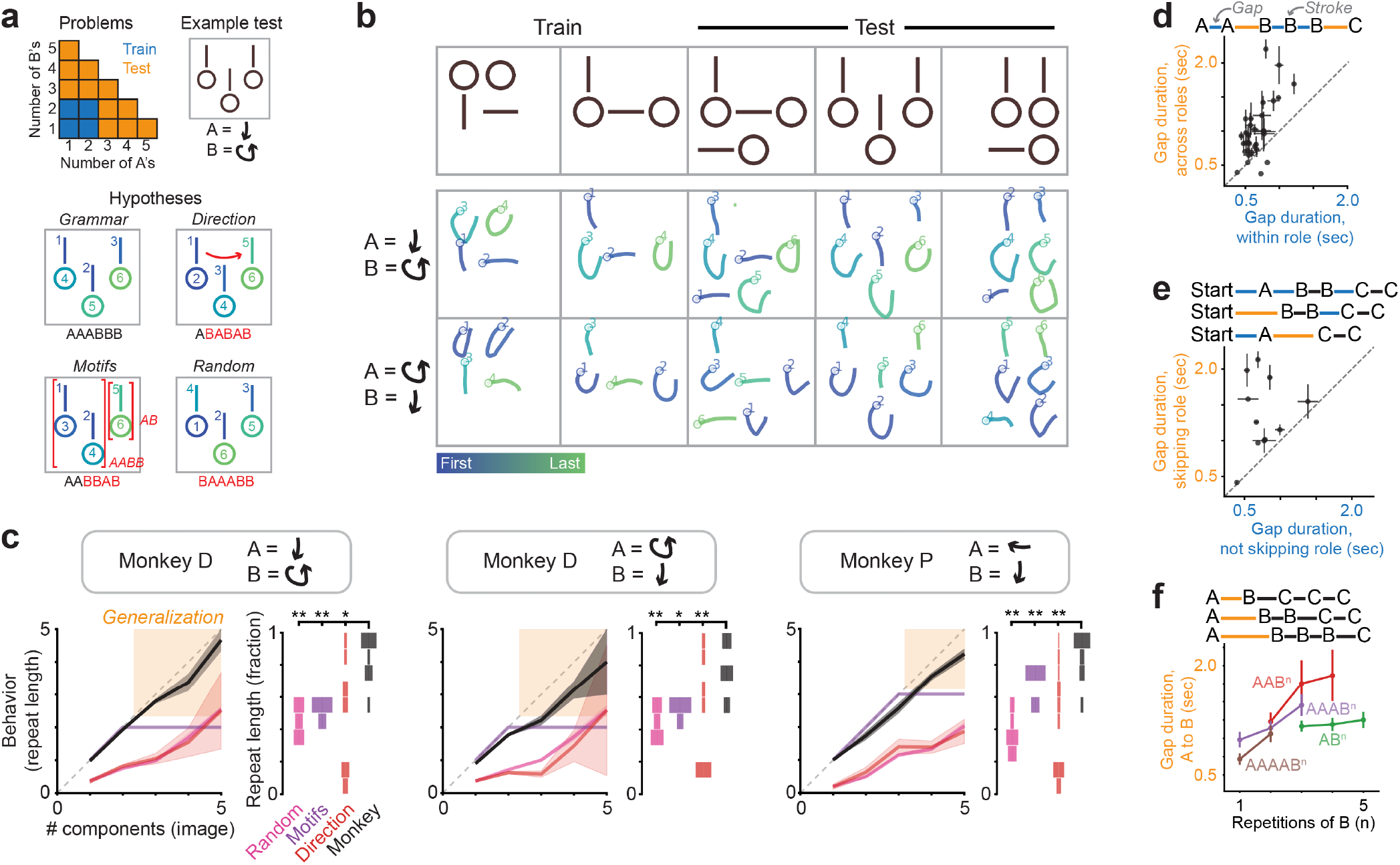
Immediate grammar-based generalization to harder problems. (a) Schematic of increase in problem complexity (more shape components) for test problems compared to training problems (top-left), along with four hypotheses for behavior on test problems (lower four panels) illustrated for an example test problem (top-right). Test problems also included new screen locations (here, the bottom row). The “grammar” hypothesis generalizes the A^n^B^m^ grammar to produce AAABBB. Considering only As produced at onset, the repeat length is three. “Direction” ignores shape content and instead sequences strokes by following a spatial direction (breaking ties in the axis perpendicular to that direction at random), which we modeled as the directional bias measured in behavior from training problems (Methods). The resulting sequence ABABAB has repeat length of one. “Motifs” reuses specific learned sequences (A^1^B^1^, A^1^B^2^, A^2^B^1^, and A^2^B^2^), resulting in chaining of AABB and AB to produce AABBAB, which has repeat length of two. “Random” samples uniformly from all possible sequences. The example random sample BAAABB has repeat length of zero. Test problems were interleaved with training problems after curriculum completion (Methods). (b) Example behavior for two training and three test problems, for the same monkey in two experiments with different shape-role associations (rows). The “line” shape includes both horizontal and vertical lines. For the three test problems, drawings depict the first attempt. (c) Summary of behavior, for two experiments for monkey D and one experiment for monkey P. Within each experiment, the left panel shows quantification of repeat length for role A at trial onset (y-axis) compared to the number of image components for role A (x-axis). We focus on A, rather than B, as it is a stronger test of grammar understanding (explained in Fig. 1i). “Monkey” represents behavior, and the other curves represent predictions for the three alternative hypotheses. Within each experiment, the right panel shows histograms of repeat length as a fraction of the number of image components, plotting probability mass over trials for the hardest test problems (four or five components). *, p < 0.05; **, p < 0.005, signed rank test, where datapoints are trials. (d) Gap duration reflects the hierarchical structure of the grammar. Top: schematic showing predicted differences in gap duration within a role (blue) and across roles (orange), for the example sequence AABBBC. Bottom: Quantification of mean gap duration for transitions across roles (y-axis) vs. within roles (x-axis). Data for monkey D (each datapoint is a unique sequence context, with one value each for transition across (e.g., AB) and within (e.g., AA) role; Methods). For monkey P, see **Extended Data Fig. 2a**. For statistical analyses that control for gap distance, see **Extended Data Fig. 2b,c**. (e) Gap duration for skipped roles reflects roles being considered in order (A to B to C). Top: schematic of predicted effect on gap duration of skipping an expected role, either skipping A to go directly from trial start to B (middle sequence) or skipping B to go directly from A to C (bottom sequence). Bottom: Quantification of mean gap duration for transitions skipping a role (y-axis) vs. not skipping the role (x-axis). Data for monkey D (each datapoint is a unique sequence context, with one value each for transition that skips a role (e.g., START-B) and does not skip a role (e.g., START-A)). For monkey P, see **Extended Data Fig. 2d**. For statistical analyses that control for gap distance, see **Extended Data Fig. 2e,f**. (f) Gap duration reflects the length of the upcoming role repeat. Top: schematic of predicted effect on A-B gap duration of increasing the repeat length of B. Bottom: mean gap duration preceding the first B (y-axis) as a function of repeat length of B (x-axis) plotted separately for different sequential contexts (colors) defined by the sequence preceding B and total sequence length. For example, the context AAB^n^ includes the sequences AABBCC, AABBBC, and AABBBB (n = 4 contexts in a single experiment for monkey D; insufficient data for monkey P). For controls, including statistical analyses that control for gap distance, see **Extended Data Fig. 2g-i.**

Monkeys progressed through a curriculum of training problems that increased in difficulty in two parallel ways: decreasing stroke-by-stroke instruction (**Fig. 1f**) and increasing problem complexity (**Fig. 1g**). This is shown for the example experiment (**Fig. 1b-k**). Stroke-by-stroke instruction refers to visual cueing of the order in which image components should be drawn (**Fig. 1f**). Over trials, instruction was reduced until it was eliminated, at which point sequencing was internally guided (**Fig. 1f**). Problem complexity, defined as the number of components in the image, started out low (two), and increased over training (up to four in this experiment, with a maximum of two lines and two circles; **Fig. 1g**). Progression through curriculum stages was dependent on performance (Methods). To motivate subjects to learn generalizable grammars rather than image-specific solutions, a large variety of images were presented (examples in **Fig. 1g**, quantification in **Fig. 1h**).

Monkeys successfully completed the training curriculum. We quantified behavior using repeat length, or the number of components drawn in repetition for a given shape (“behavior”, **Fig. 1i, j**), and compared this value to the number of image components for that shape (blue and orange, **Fig. 1i, j**), with perfect performance indicated by equality of repeat length to number of components (1 for blue, 2 for orange). Over the three-day experiment, performance improved starting on the first day, and by the end of the second day was close to perfect even with no guidance (**Fig. 1i-k**). This was true even though problems were highly variable (six examples in **Fig. 1g**; quantification in **Fig. 1h)**.

### Immediate grammar-based generalization to harder problems

To assess whether trained monkeys understood the A^n^B^m^ grammar, we tested them on problems which were harder than those encountered during training and designed to distinguish different candidate strategies (**Fig. 2a**, “hypotheses”). For the example experiment, test problems contained up to five circles and five lines, while training problems had up to two circles and two lines (**Fig. 2a**, top-left). If monkeys generalize the grammar A^n^B^m^, then they should produce correct repeat lengths for test problems, even if these lengths exceed those practiced during training. For the example problem in **Fig. 2a** with three As and three Bs, grammar use would produce a sequence (AAABBB) that repeats A and B three times each (**Fig. 2a**, “Grammar”), longer than the maximum trained repeat length of two. We also considered three alternative strategies on test problems. First, subjects may ignore the shape content and instead follow a spatial direction rule, such as producing strokes from left to right (**Fig. 2a**, “Direction”). Second, subjects may have learned the specific motifs A^1^B^1^, A^1^B^2^, A^2^B^1^, and A^2^B^2^, which would be indistinguishable from the A^n^B^m^ grammar on training problems, but would predict shorter repeat lengths on test problems. For the example problem (**Fig. 2a**, “Motifs”), this strategy would chain A^2^B^2^ and A^1^B^1^ into the sequence AABBAB, which has repeat length of two. A third alternative possibility is that test problems are perceived to be so different from training problems that the subjects have no consistent strategy, which we model as uniform sampling from all possible sequences (**Fig. 2a**, “Random”).

Subjects solved test problems by repeating all As and then all Bs, even for images with more As and Bs than in training, consistent with the grammar A^n^B^m^ (examples in **Fig. 2b**). Across subjects and experiments, we found that repeat lengths in drawings matched the number of shape components in the image (**Fig. 2c**, behavior close to identity line), including for test problems requiring longer repeats (**Fig. 2c**, “generalization”). Behavior was not consistent with the three alternative hypotheses (**Fig. 2a**), because repeats were longer than predicted by those hypotheses (**Fig. 2c**, “Monkey” is higher than “Random”, “Motifs”, and “Direction”). Success on test problems was unlikely due to additional learning on these problems, because grammar use was evident even for the first attempts (**Fig. 2c**), and reward for test problems was provided for image completion regardless of whether behavior was consistent with the grammar (Methods). In summary, subjects performed the shapes in order, repeating all cases of a shape before moving onto the next, and did so even on the first attempt for harder test problems, consistent with generalizing the grammar A^n^B^m^.

We found that monkeys could learn new assignments of shapes to the roles A and B in different experiments; the same subject drew the same test image differently depending on the current shape set (**Fig. 2b**, compare the middle to bottom row; **Fig. 2c**, compare two experiments for monkey D).

Further evidence for grammar use came from analysis of the fine-grained structure of behavioral timing. The timing of actions within sequences has been shown to reflect cognitive processes—in particular, longer duration between successive actions (gaps) may reflect planning^72^, decision making^73^, or surprise due to violation of expectations, including of grammatical structure^74^. First, we tested for hierarchical structure, where each repeat of a given role forms a high-level unit (AAABBC consists of three units: AAA, BB, C). Consistent with this, behavioral gaps between units (e.g., A to B) were longer in duration than gaps within units (e.g., A to A) (**Fig. 2d**). Second, we tested for understanding of the order of the roles A, B, and C. Consistent with this, gap duration was longer if an expected role was skipped (**Fig. 2e**). For example, A to C in AAACC is slower than A to B in AAABC, reflecting violation of an expectation that B comes after A. Third, we looked for evidence that subjects considered the entire repeat length of a role before initializing, possibly reflecting planning^72,75^ or visual search for shape components^76^. Consistent with this, gap duration before a role repeat was longer if the repeat was longer (e.g., A to B is slower for ABBB vs. ABBC) (**Fig. 2f**). These three gap duration effects were robust across experiments and not explained by distance traveled during the gap (**Extended Data Fig. 2**). Thus, generalization (**Figs. 2a-c**) and the microstructure of motor timing (**Figs. 2d-f**) provide distinct lines of evidence for understanding of action grammar.

### Identifying neural activity encoding grammatical structure

We recorded from neurons in eight frontal cortical areas previously implicated in rule-based behavior or action sequencing. Areas prominently associated with rules included dorsolateral and ventrolateral prefrontal cortex (dlPFC and vlPFC)^49,77–80^, dorsal and ventral premotor cortex (PMd and PMv)^50,81–83^, and the frontal pole (FP)^84^. Areas prominently associated with sequencing included vlPFC and dlPFC^49,52,54,58,85–87^, PMd^50,88^, the supplementary motor area (SMA)^53,89,90^, and preSMA^53,55^. PMv is also notable because its encoding of action symbols^5^ raises the possibility of contributing to grammar-based sequencing of action symbols. We also recorded from primary motor cortex (M1), an area important for motor control^47^, which has been implicated in sequencing^91,92^. Recordings were performed using chronically implanted multi-electrode arrays (sixteen 32-channel arrays; **Figs. 3a, b; Extended Data Fig. 3**). Recording from these areas at the same time allowed us to perform the first direct comparison of their activity in the same task. We first assessed the extent to which activity during drawing encodes the roles A, B, and C (**Fig. 3**) and does so in a manner abstract over specific shape content (**Fig. 4**).

**Figure 3.**
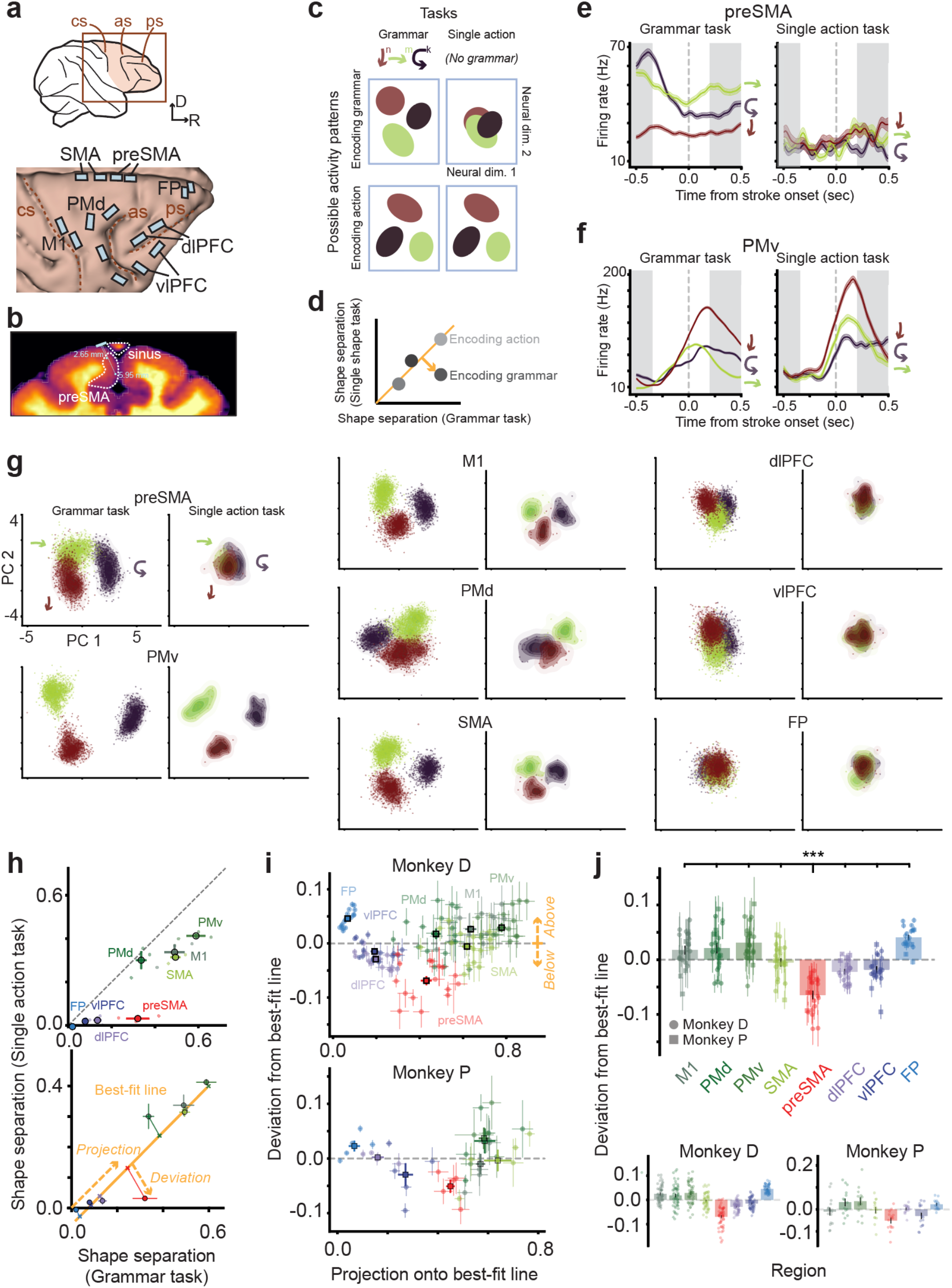
Encoding of grammar-specific roles A, B, and C in preSMA. (a) Recording sites, targeted to right frontal cortex contralateral to the drawing hand (top), showing 32-channel floating microelectrode arrays (bottom) to scale on 3D rendering of brain (monkey P). cs, central sulcus; as, arcuate sulcus; ps, principal sulcus. R, rostral. D, dorsal. (b) Coronal MRI section depicting SMA and preSMA arrays angled to avoid the superior sagittal sinus and target the medial wall, for the most caudal preSMA array. Arrays were implanted ∼2 mm lateral to the midline, and angled medially. The two lines represent the shortest (2.65 mm) and longest (5.95 mm) electrodes. This location and angle is our best estimate based on the surgical plan, stereotactic coordinates, and intra-operative photographs. The other SMA and preSMA arrays were angled similarly. D, dorsal. L, lateral. (c) Schematic of predicted neural activity encoding roles for hypothetical populations (rows) that either encode roles in the grammar (“Encoding grammar”) or encode actions regardless of the grammar (“Encoding action”), across two tasks (columns) that either require the use of the grammar (“Grammar”) or not (“Single action”). Population activity is projected onto a two-dimensional subspace (same between columns) during drawing of *A* (maroon ellipse), *B* (green), and *C* (black). (d) Schematic of shape separation quantification indicative of a region that encodes grammar-specific roles, quantifying the schematic in panel c. This plots the magnitude of separation in neural space between different shapes (shape separation; see below) during the Single action task (y-axis) and Grammar task (x-axis) for four hypothetical brain regions (grey points). Regions that encode action should have similar shape separation between tasks (falling along the orange line) and regions that encode grammar-specific roles should have high shape separation during the Grammar task and low shape separation during the Single action task (deviating towards the bottom right). (e) An example unit in preSMA, showing mean activity for each role (colors) during the Grammar or Single action task. White background indicates the time window aligned to stroke onset used for subsequent analyses (−0.35 to 0.2 sec). (f) An example unit in PMv, recorded concurrently and plotted analogously to panel e. (g) Population activity for eight simultaneously recorded brain regions, each projected to two dimensions using PCA (Methods), plotting for each role a kernel density estimate (KDE; filled contours) and individual stroke datapoints, in the time window using data −0.35 to 0.2 sec relative to stroke onset. For each region, Grammar and Single-action task data are combined before PCA and then data from both tasks are projected into the same subspace. (h) Shape separation for an example experimental session, comparing the Grammar task (x-axis) and the Single action task (y-axis). The top panel shows all datapoints (one point for each shape). The bottom panel illustrates the approach to quantify each region’s deviation from the overall action encoding axis. This axis was estimated using ordinary least squares regression. We quantified each region’s projection and orthogonal deviation relative to that axis. (i) Overview of orthogonal deviation (y-axis) versus projection (x-axis). Each circle represents one session (mean +/- SEM over shape kinds) and each square represents the mean +/- SEM over sessions. (j) Summary of orthogonal deviation across brain regions. In the top panel each point represents one session (mean +/- SEM over shape kinds) and bars represent the mean +/- SEM over sessions (combined across animals). The lower two plots split the data by animal; circle points represent shape types pooled across sessions, and bars represent mean +/- SEM over shape types. ***, p<0.0005, preSMA vs. each other region, linear mixed effects model with fixed effect of brain region and random effects for animal and session. Datapoints were shape kinds pooled across sessions.

**Figure 4.**
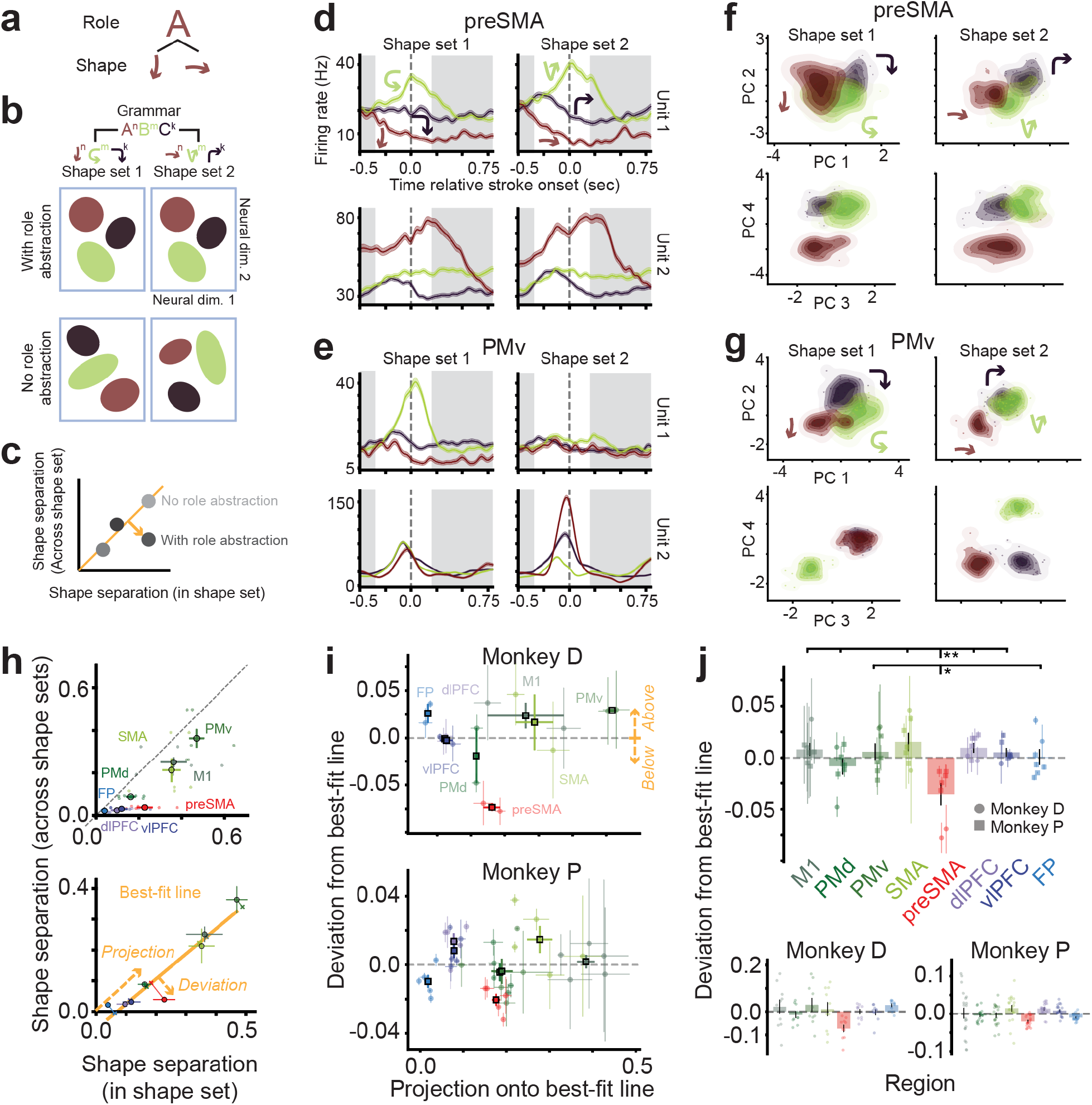
Role representations exhibit abstraction over specific shapes. (a) Schematic of role abstraction; the role A can be filled by either of these two shapes. (b) Schematic of predicted neural activity reflecting role abstraction for hypothetical populations (rows) that either exhibit abstraction (“With role abstraction”) or encode specific actions (“No role abstraction”), in a task that requires switching on a trial-by-trial basis between two shape sets (columns). (c) Schematic of shape separation quantification indicative of role abstraction, showing neural separation between shapes computed either within each shape set and averaged across sets (x-axis) or between the two shape sets within each role and then averaged across roles (y-axis), for four hypothetical brain regions (grey points). Regions exhibiting role abstraction have high separation within shape sets and low separation across sets, thus deviating towards the bottom right. (d) Two example units (rows) in preSMA, showing mean stroke-aligned activity for each of the three shapes in the two shape sets (columns). White background indicates the time window used for subsequent analyses (−0.35 to 0.2 sec). (e) Two example units (rows) in PMv, recorded concurrently and plotted analogously to panel d. (f) Population activity for preSMA, projected to two 2D subspaces (rows) spanning the first four principal components, split by shape set (columns). Each plot shows a KDE and individual datapoints for each shape (colored by role) in a given shape set, and for a specific screen location, using data in the time window of −0.35 to 0.2 sec relative to stroke onset. For each region, data are combined between shape sets before PCA and then data from both shape sets are projected onto the same subspace. (g) Population activity for PMv, recorded concurrently and plotted analogously to panel f. For the six brain regions other than preSMA and PMv, see **Extended Data Fig. 4**. (h) Shape separation for an example experimental session. The top panel shows all datapoints (one point for each shape). The bottom panel illustrates the approach to quantify each region’s deviation from the overall action encoding axis. We quantified each region’s projection and orthogonal deviation relative to a best-fit axis identified using ordinary least squares regression across regions. (i) Overview of orthogonal deviation (y-axis) versus projection (x-axis) relative to each session’s action encoding axis. Each circle represents one session (mean +/- SEM over shape kinds) and each square represents the mean +/- SEM over sessions. (j) Summary of orthogonal deviation across brain regions. In the top panel each point represents one session (mean +/- SEM over shape kinds) and bars represent the mean +/- SEM over sessions (combined across animals). The lower two plots split the data by animal; circle points represent shape types pooled across sessions, and bars represent mean +/- SEM over shape types. *, p < 0.05; **, p<0.005, preSMA vs. each other region, linear mixed effects model with fixed effect of brain region and random effects for animal and session. Datapoints were shape kinds pooled across sessions.

### Representation of grammar-specific roles A, B, and C in preSMA

Encoding of roles during drawing should be evident as distinct activity states during production of A, B, and C. Critically, if these states encode roles, they should correspond to internal representations that are active during grammar-based behavior and not strongly driven just by image or motor properties of the shape. We thus expected that activity representing roles should be stronger in a task that requires producing a sequence using the grammar (**Fig. 3c**, “Grammar task”) compared to a control task that requires drawing one stroke per trial and therefore does not require the grammar (**Fig. 3c**, “Single action task”). This expectation for a hypothetical population encoding roles is illustrated in a schematic of neural activity (**Fig. 3c**, “Encoding grammar”). At the other extreme, a population encoding individual actions rather than grammar should show similar activity between Grammar and Single action tasks (**Fig. 3c**, “Encoding action”). In a quantification **(Fig. 3d**), a population reflecting roles is predicted to exhibit high difference in activity across shapes during the Grammar task (**Fig. 3d**, x-axis) and low difference during the Single action task (**Fig. 3d**, y-axis).

Out of all areas, we found that preSMA most strongly exhibited role encoding, evident as shape encoding that was strong during the Grammar task and weak during the Single action task. This is evident for an example preSMA unit (**Fig. 3e**); its stroke-aligned activity differed strongly by shape in the Grammar task (**Fig. 3e**, left), but not in the Single action task (**Fig. 3e**, right). An example unit in PMv also encoded shapes during the Grammar task (**Fig. 3f**, left). However, in contrast to the preSMA unit, this encoding was also strong during the Single action task (**Fig. 3f**, right), consistent with encoding not of roles, but of action symbols that generalize over sequential contexts^5^. These encoding properties were also evident at the population level. Two-dimensional linear projections of preSMA population activity revealed separate shape-specific states during the Grammar task, which diminished for the Single action task (**Fig. 3g**, “preSMA”). In contrast, the other seven brain regions exhibited either strong shape separation in both tasks (PMv, PMd, SMA, M1; **Fig. 3g**) or weak separation in both tasks (dlPFC, vlPFC, FP; **Fig. 3g**).

We quantified role encoding in the following manner. First, to quantify the extent to which each region’s population activity differed by shape, we devised a “shape separation” metric, which is the average pairwise Euclidean distance between all across-shape trials, subtracting the average within-shape Euclidean distance, normalized between 0 (no difference) and 1 (a ceiling, defined as the 98th percentile of pairwise Euclidean distances). This metric was computed in a manner that controlled for potentially confounding behavioral factors, including the stroke’s location and preceding movement direction (Methods). In analysis of shape separation for all regions, preSMA had relatively high shape separation in the Grammar task (**Fig. 3h**, x-axis) and low separation in the Single action task (y-axis). From shape separation scores, we then quantified role encoding as the extent to which a given region had weaker Single-action shape separation than expected from the overall across-region relationship between Single-action shape separation and Grammar shape separation (**Fig. 3d**, length of orange arrow). For each session, we used linear regression to identify a line that captures the overall relationship between Single-action and Grammar shape separation (**Fig. 3h**, bottom, “Best-fit line”) and computed each region’s projection along that line and orthogonal deviation from that line (**Fig. 3h**, bottom). In the resulting scatterplot of deviation versus projection (**Fig. 3i**), greater projection (higher along the x-axis) reflects stronger action encoding and greater negative deviation (lower along the y-axis) reflects stronger role encoding.

Projection scores increased from prefrontal and frontal polar areas (FP, dlPFC, vlPFC), to preSMA, to premotor and motor areas (PMv, PMd, SMA, M1) (**Fig. 3i**), reflecting a known increase in movement-related activity along the anterior-to-posterior gradient of frontal cortex^93^. Deviation scores did not follow such a gradient. Instead, they were maximally negative for preSMA (**Fig. 3i**). Direct comparison to every other region revealed that preSMA exhibited the most strongly negative deviation scores (**Fig. 3j**). This negative deviation indicates encoding of grammar-specific role in preSMA. Thus, we identified activity that encodes the roles A, B, and C as distinct activity states preferentially active during grammar use, and this encoding was strongest in preSMA.

### Role representations in preSMA exhibit abstraction over specific shapes

High-level cognition has been hypothesized to involve role representations that are abstract over specific objects, as in theories involving dynamic role-filler binding^35,36^ and algebraic sequences^4,10,13^ (**Fig. 4a**). For example, in language, understanding the roles of “subject”, “verb”, and “object” allows generalizing to any sentence with that structure, such as “John loves Mary” and “Mary hates John”. To assess whether neural activity encodes the roles A, B, and C in a manner that is abstract across specific shapes, we trained macaques to switch between two shape sets that share the same abstract A^n^B^m^C^k^ grammar (**Fig. 4b**, top). Switching occurred on randomly interleaved trials and the two sets had different shapes. If role representations exhibit abstraction, then activity should be similar across shape sets (**Fig. 4b**, “With role abstraction”). In contrast, if activity encodes the specific shapes, then activity should differ (**Fig. 4b**, “No role abstraction”). Accordingly, role abstraction predicts high activity difference between shapes within each shape set (**Fig. 4c**, x-axis) and low difference between shapes with the same role but in different shape sets (**Fig. 4c**, y-axis).

We found evidence for role abstraction, strongest in preSMA. Example preSMA units (**Fig. 4d**) exhibited similar activity patterns between shape sets 1 and 2 (columns), consistent with abstraction, while example PMv units (**Fig. 4e**) had different activity between shape sets; for example, one PMv unit fired strongest for the “C” shape (unit 1, shape set 1, green) and the other fired strongest for the rightwards line (unit 2, shape set 2, red). These effects were evident also in visualizations of population activity, in which the first four principal components of preSMA activity were similar between shape sets 1 and 2 (**Fig. 4f**). In contrast, other brain regions exhibited either strong differences between shape sets (PMv in **Fig. 4g**; M1, PMd, SMA in **Extended Data Fig. 4**) or lacked strong shape separation even within shape sets (dlPFC, vlPFC, FP; **Extended Data Fig. 4**).

We quantified abstraction in population neural activity in the following manner and analogous to role encoding. First, we computed two shape-separation metrics for each area: shape separation within each shape set (**Fig. 4h**, x-axis) and shape separation within the same role but across shape sets (**Fig. 4h**, y-axis). We quantified abstraction as the extent to which a given region has weaker across-set shape separation than expected from the overall relationship between across-set separation and within-set separation (**Fig. 4c**, length of orange arrow). For each session, we used linear regression to identify a line that captured the overall relationship between across-set and within-set shape separation (**Fig. 4h**) and computed each region’s projection and deviation relative to that line, with greater negative deviation indicating stronger abstraction. We found that deviation scores were most negative for preSMA compared to every other region (**Fig. 4i, j**), indicating role abstraction in preSMA. Together, these results identify neural activity that encodes roles in a manner abstracted over specific shapes, with strongest levels of abstraction in preSMA.

### Representation of repetition ordinal position in preSMA

Neural activity representing stroke-by-stroke progression (ordinal position) within role repeats (**Fig. 5a**, “Encoding ordinality”) would be consistent with a strategy that internally tracks progression, possibly with respect to a goal defined by a target number of repetitions. On the other hand, because the target image stays visible during drawing (**Fig. 1e**), this task could be solved without an internal representation of ordinal position; instead, the subject could visually compare the state of the current drawing (represented on screen as visible “ink”) to the target image.

**Figure 5.**
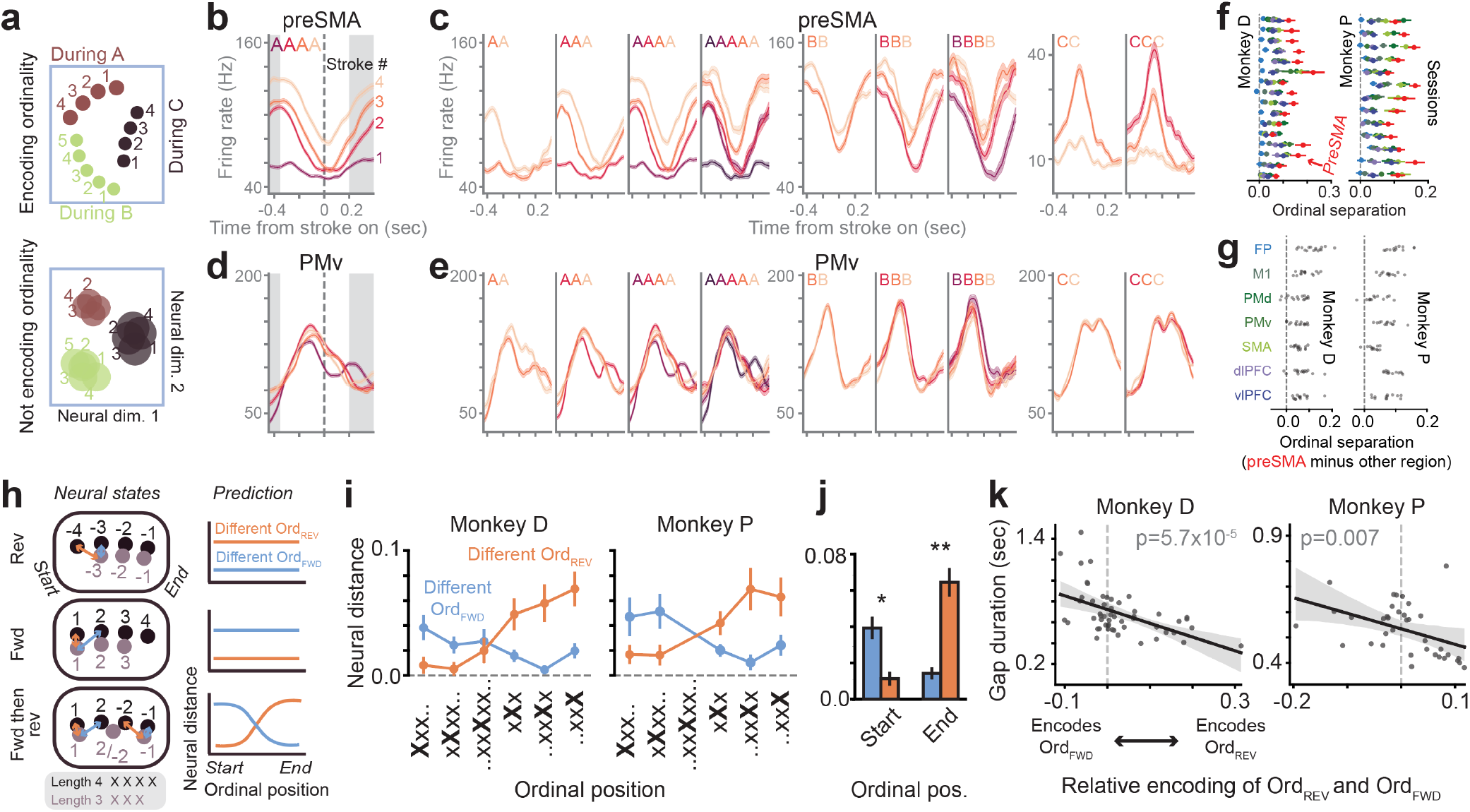
Encoding of repetition ordinal position in preSMA. (a) Schematic of ordinal position encoding depicting predicted neural activity for hypothetical populations (rows) that either encode ordinal position within roles (“Encoding ordinality”) or do not (“Not encoding ordinality”). (b) An example unit in preSMA, showing mean stroke-aligned activity for each ordinal position (numbered, counting up from repeat onset) during length-4 repeats of A. White background indicates the time window used for subsequent analyses (−0.35 to 0.2 sec). (c) Example unit activity in preSMA. Each subpanel is a subset of the data during a specific length repeat of a specific role. Mean activity is shown for each ordinal position. The same unit from panel b is plotted for *A* and *B* repeats and a different unit for *C* repeats. (d) An example unit in PMv, recorded concurrently and plotted analogously to panel b. (e) Example unit activity in PMv recorded concurrently and plotted analogously to panel c. This is the same unit from panel d. (f) Summary of ordinal separation, one mean value for each brain area (colors; see colormap in panel g) in each session. (g) Summary of ordinal separation, showing the difference between mean encoding in preSMA vs. each other area; n = 13 sessions (monkey P), 16 (D). Statistics: p < 0.001, for each preSMA vs. other area comparison, for both monkeys P and D. (h) Schematics depicting comparison of different-length repeats—here, 3 (black) and 4 (grey)—to test whether activity represents ordinal position from repeat end (ord_REV_) (top), ordinal position from repeat start (ord_FWD_) (middle), or a hybrid scenario representing ord_FWD_ at repeat start and then ord_REV_ at repeat end. Arrows (in “Neural states”) represent neural distance between states with different ord_FWD_ but same ord_REV_ (blue) or different ord_REV_ but same ord_FWD_ (orange). These three scenarios make different predictions for the relationship between neural distances and the ordinal position at which the distances are computed (“Prediction”). (i) Comparison of strength of encoding of ord_FWD_ and ord_REV_ as a function of ordinal position along a repeat. For each datapoint—a specific ordinal position, repeat length, and role (x-axis)—we computed its neural distance (mean +/- SEM) to ordinal positions for a repeat of the same role but different length, such that each datapoint contributed one value each for “different ord_FWD_ and same ord_REV_” (blue) and “different ord_REV_ and same ord_FWD_” (orange). X-axis labels represent ordinal position (indicated by location of capital X in sequence) and repeat length, with “..” indicating the possibility of more items of “x”. (j) Summary of forward (blue) and reverse (orange) ordinal encoding, computed either at the start or end of repeats. “Start” includes positions with ord_FWD_ < 3 and ord_REV_ < −2 (Xxx.. and xXxx.. in panel i) and “end” includes positions with ord_FWD_ > 2 and ord_REV_ > −3 (..xxXx and ..xxX in panel i). Data are combined across two subjects. *, p < 0.05; **, p < 0.005, comparing “different ord_FWD_” to “different ord_REV_” using the Wilcoxon signed-rank test, performed independently for each animal. (k) Relationship between gap duration and relative encoding of ord_FWD_ and ord_REV_, computed as d_REV_ – d_FWD_, where d_REV_ is the neural distance between states differing in ord_REV_, and d_FWD_ is the distance between states differing in ord_FWD_. This shows results from an ordinary least squares regression fit (95% bootstrapped CI). Each datapoint represents a specific role from an experimental session, with gap duration averaged over gaps, and ordinal encoding computed separately for each ordinal position and then averaged.

Consistent with monkeys internally tracking progression towards completion of a role repeat, we found that preSMA activity strongly differed by ordinal position within repeats. An example preSMA unit’s activity differed systematically by ordinal position within length-4 repeats of A (AAAA, **Fig. 5b**). Activity differed by ordinal position in a similar manner (increasing in firing rate) for different length repeats of A (**Fig. 5c**, AA to AAAAA). Its activity also differed by ordinal position for repeats of B, but in an opposite manner (decreasing in firing rate) (**Fig. 5c**, BB to BBBB). In contrast, activity in PMv did not strongly differ by ordinal position. This is evident in the example PMv unit for AAAA (**Fig. 5d**) and across different length repeats and roles (**Fig. 5e**). These effects—ordinal encoding that was strong in preSMA and weak in PMv—were also evident in population activity (**Extended Data Fig. 5**). We quantified ordinal separation (the mean pairwise neural distance between all ordinal positions, computed separately for each role and averaged) and found that it was strongest in preSMA compared to each other area (**Fig. 5f, g**). Thus, preSMA activity differs by ordinal position during repeats.

To determine whether ordinal position encoding reflected the number of remaining items in a repeat, we compared neural activity across trials with different length repeats (**Fig. 5h-k**). This approach evaluated whether activity encodes ordinal position relative to the end of a repeat (“number of items remaining”, termed *ord_REV_*; schematized in **Fig. 5h**, top) or relative to the start of a repeat (“number of items completed”, *ord_FWD_*; **Fig. 5h**, middle). A third possibility is that activity is “anchored” at repeat boundaries, such that it represents *ord_FWD_* at repeat onset and switches to *ord_REV_* at repeat offset (**Fig. 5h**, bottom). These three possibilities make different predictions for the neural distances between ordinal positions from different length repeats (**Fig. 5h**, right column). For example, encoding of ord_REV_ (**Fig. 5h**, top) predicts that activity for Xxx (the first stroke in a length-3 sequence of role X) differs from Xxxx activity (long orange arrow) and is similar to xXxx activity (short blue arrow). In contrast, encoding of ord_FWD_ (**Fig. 5h**, middle) predicts that Xxx is similar to Xxxx and different from xXxx. Taking this approach, we found that preSMA activity was consistent with the third, “anchored” model (data in **Fig. 5i**; prediction in **Fig. 5h**, bottom), encoding ord_FWD_ at repeat onset and then changing to encode ord_REV_ with around three strokes remaining (**Fig. 5i, j**). This result suggests that subjects internally track progression within repeats in a dynamic manner, “counting up” at the start and then “counting down” towards the end.

This finding of activity encoding ord_REV_ towards the end of repeats suggests a representation of the number of remaining repetitions in a plan for upcoming actions. Having a plan would minimize the need for further decision making or evaluation of the partially completed image before each stroke, which would predict shorter gap durations between strokes. Consistent with this, sequences with stronger encoding of ord_REV_ also had shorter gap duration (**Fig. 5k**). Thus, representation of ord_REV_ may be a neural signature of a plan for the remaining actions within a repeat.

### Population geometry of grammar structure in preSMA

We have found that population activity in preSMA reflects abstract grammar-specific roles (**Fig. 3, 4**) and ordinal position within each role (**Fig. 5**). We next assessed population geometry—the overall relational structure defined by the distances and angles between activity states representing distinct roles and ordinal positions^80^. We considered three key relational properties of neural activity. First, ordinal positions within roles may be organized along linear axes or may be unstructured (**Fig. 6a**, “Ordinal”). The existence of linear axes predicts high alignment between vectors taken between successive ordinal positions (**Fig. 6a**, “Ordinal”, cosine similarity of 1), while an unstructured organization predicts low alignment (**Fig. 6a**, “Ordinal”, cosine similarity of 0). Second, ordinal linear axes, if they exist, may be parallel across roles, predicting high alignment between ordinal-axis vectors across roles (**Fig. 6a**, “Ordinal (different role)”, similarity of 1). Third, roles may be organized along a linear axis (**Fig. 6a**, “Role”, similarity of 1) or unstructured (**Fig. 6a**, “Role”, similarity of 0). Together, different combinations of these three relational properties define multiple ways that roles and ordinal positions can be inter-related (**Extended Data Fig. 6**).

**Figure 6.**
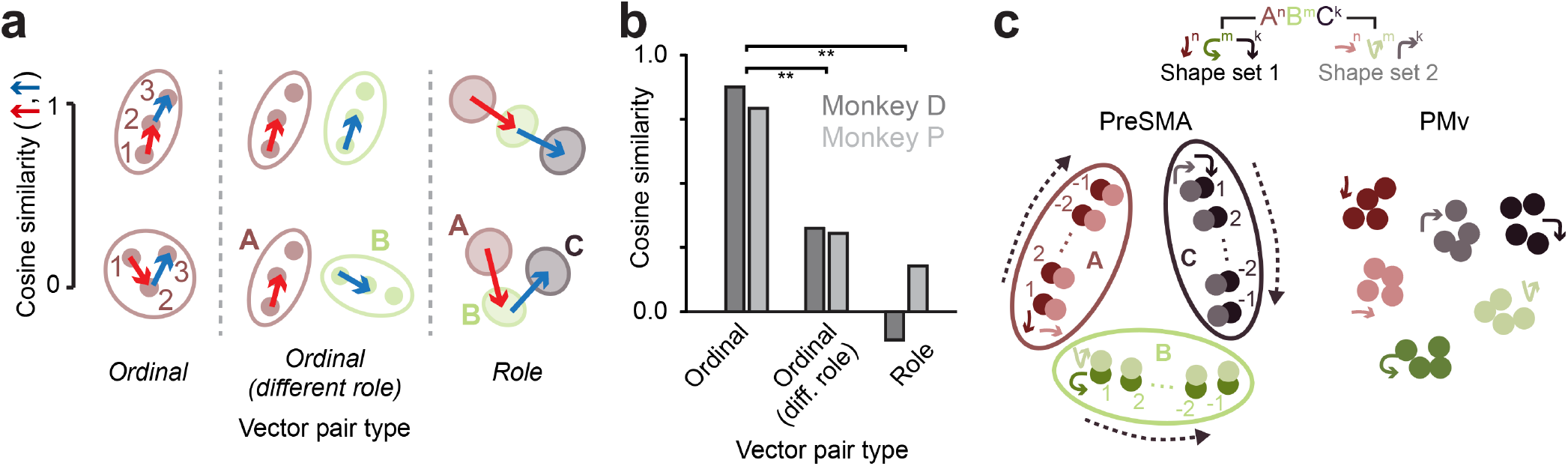
Population geometry of grammar structure in preSMA. (a) Schematic of three relational properties we assessed in population neural activity. Each property falls along a continuum of increasingly linear or collinear organization (vertical axis), quantified as the angle (cosine similarity) for a specific type of vector pair (horizontal axis). Each vector type is taken between a pair of specific neural activity states (blue and red arrows). Roles (A, B, C) are depicted in three colors and ordinal positions (1, 2, 3) are depicted as small, filled circles. (b) Summary of angles for each type of vector pair in neural population data. Cosine similarity is computed using trial-averaged dot products (Methods); **, p < 0.005, permutation test, performed by shuffling vector-pair labels at the level of dot products (Methods), testing the hypothesis that values for “ordinal” vector pair are different than values for the other vector pairs. For each animal, the average is over all sessions that have data across all vector pairs. (c) Schematic summarizing the preSMA population geometry inferred from the results of this study. The schematic for PMv is based on both the current and a previous study^5^.

Analysis of pairwise angles between state-to-state vectors (**Fig. 6b**) revealed the presence of linear axes encoding ordinal positions (high “Ordinal” cosine similarity). These axes were close to orthogonal across roles (**Fig. 6b**, low “Ordinal (diff. role)” similarity). This analysis also revealed largely unstructured role representations (**Fig. 6b**, low “Role” similarity). This analysis, combined with the results from **Figs. 3-5**, reveals a particular representational geometry of grammar structure in preSMA (**Fig. 6c**): each role is represented by a linear axis (A, B, C) that encodes ordinal progression anchored to the first and last strokes (1, 2, …, −2, −1). These role axes are abstract over specific shapes (two shapes shown per role) and are largely orthogonal across roles.

## Discussion

### Identification of grammar structure independent of action content

Whether and how the brain represents abstract grammar structure independently of specific objects, actions, or events—thereby allowing structure-level operations to generalize across arbitrary content—is a central question in the study of cognition^7,11,13,34–36,94^. We identified a neural population in preSMA encoding action grammar structure (summarized in **Fig. 6c**), distinct from a neural population that encodes action content, in PMv^5^ (**Fig. 6c**). In macaques solving drawing-like problems requiring action sequences using the grammar A^n^B^m^C^k^ (**Figs. 1, 2**), preSMA activity encodes the abstract roles A, B and C as distinct grammar-specific states (**Figs. 3, 4, 6**) and encodes ordinal positions along linear axes that are largely orthogonal across roles (**Fig. 5, 6**). Together, these findings identify neural activity representing abstract structure of a grammar, link that activity to a specific brain region (preSMA), and characterize a relational structure encoded by that activity’s population geometry.

Grammar structure and action content were separated in independent codes and into distinct populations. They used independent codes, in that role representations (in preSMA) were minimally influenced by the shape filling the role (**Fig. 4**), and shape representations (in PMv) were minimally influenced by role (**Fig. 4**; consistent with prior evidence that PMv encodes action symbols invariant to sequence context and motor kinematics^5^). Thus, during drawing, distinct codes simultaneously represent structure (e.g., role A) and content (shape assigned to role A) (**Fig. 6c**). A second implication is that grammar and action are preferentially encoded in distinct populations, supported by our findings of differences between preSMA and PMv. Future studies may clarify whether and how grammar structure and action content are bound^35,36,40^ into a unified representation through interactions between the interconnected^95,96^ populations in preSMA and PMv.

Evidence for the representation of abstract roles in macaques deepens our understanding of the neural basis of high-level cognitive abilities in primates, and, potentially in turn, the neural and evolutionary basis of human cognition. Role representations that abstract over content have been hypothesized in human cognition, including in language^7,94,97^, relational and logical reasoning^7,35,41^, concept learning^40^, navigation^21^, sequence learning^10^, and action^22^, and can emerge in artificial neural networks optimized to perform cognitive tasks^37,98^. Yet, while non-human animals can readily grasp ordinal structure^10,99^, whether they understand abstract roles has remained a subject of debate^4,10–12,21,100^. Our study’s finding of abstract role representations raises the possibility that such representations may be a more general feature of primate cognition than previously recognized; moreover, they may relate to prior findings in primates, including neural correlates of abstract ordinal position during sequencing^52,59,61,90,101–104^, neural correlates of categories^54,77,105^, and the behavioral capacity to learn the order of categories^106^. More speculatively, the neural substrates of role representations in this study may relate to the neural and evolutionary bases of human cognition.

### A neural representational basis of hierarchical grammar structure

The identified representational geometry (summarized in **Fig. 6c**) provides a more direct view of the internal representation of this grammar than behavior. Behavioral findings—successful generalization (**Fig. 2a-c**) and the microstructure of gap durations (**Fig. 2d-f**)—indicated a hierarchical grammatical structure consisting of a high-level understanding of the roles and their order (A then B then C) and a low-level understanding to repeat each role until all its instances are drawn. Complementing these findings with recordings revealed three key properties of the underlying structure of this grammar. First, in principle, this hierarchical structure in behavior could have corresponded to multiple possible neural geometries, including (1) a structure with a single “global” axis that chains together successive role repeats (**Extended Data Fig. 6**, vii), potentially related to preSMA activity that encodes global ordinal position^59,101^, or (2) a structure with two orthogonal axes^80^, one encoding role progression and another encoding role-general ordinal progression (**Extended Data Fig. 6**, vi), reflecting a general-purpose system for ordinal encoding^107^. Instead of these two possibilities, we identified a structure with role-specific near-orthogonal ordinal axes (**Fig. 6c**; **Extended Data Fig. 6**, iii). Second, repetition of a role could have corresponded to ordinal axes anchored at onset or offset of repeats (**Fig. 5h**); instead, we found that axes were anchored at both the onset and the offset of repeats (**Fig. 5h-j**)—a property that has been found in other sequencing tasks^60,70,108^ and thus may reflect a general neural coding mechanism. Third, each role could have corresponded to a distinct subpopulation of neurons (or a specific population subspace), whose activity encodes what shape is assigned to that role (i.e., conjunctive role-shape codes)^34,109^, resembling recently reported ordinal subspaces (“slots”) in sequence working memory^70,85^. Instead, we identified neurons encoding role in a manner abstract over specific shapes (**Fig. 4**). Thus, neural recordings revealed a specific internal representation of grammar structure.

The identified representation suggests mechanistic hypotheses for generalization ability. First, our finding of representations of abstract roles suggests the presence of mechanisms that flexibly associate, or bind, role representations with specific shapes across two timescales, both over gradual learning (**Fig. 2b**) and on a trial-by-trial switching basis (**Fig. 4**). Second, finding that ordinal axes are anchored at their onsets and offsets (**Fig. 5h-j**) suggests that a way to generalize across repeat lengths is to adjust the velocity of neural trajectories along ordinal axes to compress the trajectory for longer repeats and expand it for shorter ones^108,110^. Such an operation may benefit from ordinal axes being largely orthogonal across roles (**Fig. 6**), allowing axes to vary independently within individual trials. Anchoring may also reflect a particular cognitive strategy, in which macaques plan the number of remaining repetitions when there are around three strokes remaining (**Fig. 5h-k**). Thus, the identified grammar representation suggests hypotheses for mechanisms underlying grammar generalization.

### A central role for preSMA in grammar-based sequencing

Our results suggest that a function of preSMA is to represent and implement action grammars. This function accounts for, and extends, previous findings suggesting a role for preSMA in sequencing based on neural activity correlates of sequence structure^55,71^, including of ordinal position^59,90,101,111,112^, sequence context^90,113,114^, and future actions^114^. Our study differs from prior work in that it (i) identifies abstract role encoding (in combination with ordinal encoding), (ii) quantifies population geometry, revealing a single relational structure that incorporates both role and ordinal encoding, (iii) directly compares preSMA to other frontal areas (although see ref^59^), and (iv) does this in a task involving grammar-based generalization rather than explicit instruction or memory (see below). With these advances, our study demonstrates the abstract, hierarchical, distinct, and generative nature of preSMA activity, four properties which jointly support a central role for this area in grammar-based sequencing.

Grammar-based sequencing may in turn account for diverse functions associated with preSMA in humans and monkeys. These include movement-related functions^115^, including tracking time and rhythm^105,116,117^, updating plans^90,118,119^, learning new sequential procedures^53,120^, generating sequences^55^, and initiating action by oneself without using external cues^115,121,122^. Beyond movement, preSMA is also implicated in a striking array of abilities^123^, including language^124^, musical improvisation^123,125^, creative drawing^126^, handwriting^127^, mental visuo-motor rotation^128^, arithmetic^123,129^, sequencing operations^128,130^, and abstract formal reasoning^130,131^. Our study raises the possibility that many of these findings reflect a single fundamental function of preSMA, which is to represent grammar structure.

While there is substantial evidence for involvement of lateral prefrontal cortex in sequencing^49,54,58,85,87,132,133^ and using abstract rules^49,78–80^, grammar structure was surprisingly weak in that region (dlPFC and vlPFC) compared to preSMA. Two key innovations in our study facilitated the discovery of a dominant role for preSMA in grammar-based sequencing. First, we recorded from many areas previously linked to sequencing or rule use, allowing us to quantify differences across these areas in the same animals and trials. Second, our task isolated grammar-based sequencing from the need to follow instructive cues or use memorization, two methods commonly used in prior studies (see below). A fruitful direction for future studies will be to understand the interaction between preSMA, lateral prefrontal areas, and other cortical and subcortical areas, including how they coordinate to implement grammar-based sequencing, and how disruption of activity in these areas may account for disorganization of thought and action in frontal lobe disorders^134^.

### A task isolating grammar-based generalization

A key advance of our study is the establishment of an animal task to test grammar-based action sequencing, a kind of sequencing that has been understudied in prior work, which has instead focused on instructed or memorized sequences. Instructed sequences use cues that explicitly guide action order (e.g., spatial targets that light up in sequence)^47,48,51,52,85,87,111,135^ and memorized sequences involve reproduction of a learned sequence, either highly practiced^48,57,91,120,136,137^ or from working memory^54,55,88,103,104,114^. In contrast, many real-world tasks require grammar-based sequencing: they present no instruction and involve highly varied unfamiliar problems. In such cases, solutions must be inferred by applying general rules to unfamiliar problems. Accordingly, grammar-based sequencing has been a focus of behavioral studies^32,33,63,64,66,138^. We developed a task that involves no instruction (static images without ordering cues) and disincentivizes memorization, both short-term (the image stays on) and long-term (images are highly varied and can be novel). Instead, subjects inferred the correct sequence for a given image by applying a learned grammar. Critically, they immediately generalized, solving new problems on the first attempt (**Fig. 2**), indicating that this task successfully incentivizes and assesses the generative ability essential to grammars.

### A frontal cortical system for compositional generalization

The present findings suggest that preSMA and PMv are two components of a putative frontal system that supports compositional generalization. Understanding whether and how neural representations and dynamics in this system realize symbolic, program-like operations^11,40–42,49,139,140^ may help bridge long-standing symbolic and connectionist theories of cognition^2,7,12,37,43,94,109,141,142^, clarify how abstract roles and content are dynamically combined^6,34–36,40^, and generate new insights into the neural basis of intelligence.

## Methods

### Subjects and surgical procedures

Data were acquired from two adult male macaques [*Macaca mulatta*, average weights 17 kg (Monkey P) and 10 kg (Monkey D), average ages 10 years (P) and 8 years (D)]. This sample size was chosen to match the standard for neural recording studies of behaving monkeys^5^. All animal procedures complied with the NIH Guide for Care and Use of Laboratory Animals and were approved by the Institutional Animal Care and Use Committee of the Rockefeller University (protocol 24066-H).

After undergoing initial task training in their home cages, subjects underwent two surgeries, the first to implant an acrylic head implant with a headpost, and the second to implant electrode arrays, as described in detail in a previous study^5^ and described in brief here. Both surgeries followed standard protocol, including for anesthetic, aseptic, and postoperative treatment. In the first surgery, a custom-designed MR-compatible Ultem headpost was implanted, surrounded by a bone cement cranial implant, or “headcap” (Metabond, Parkell and Palacos, Heraeus), which was secured to the skull using MR-compatible ceramic screws (Rogue Research). After a six month interval, to allow bone to grow around the screws and for the subject to acclimate to performing the task during head fixation via the headpost, we performed a second surgery to implant 16 floating microelectrode arrays (32-channel FMA, Microprobes for Life Science), following standard procedures^5,143^. The craniotomy was then closed with bone cement.

Two arrays were targeted to each of multiple areas of the frontal cortex, with locations identified stereotactically, and planned using brain surface reconstructions derived from anatomical MRI scans (3D Slicer 5.6.2). Locations were selected based on their published functional and anatomical properties, anatomical sulcal landmarks, and a standard macaque brain atlas^144^, as detailed in a previous study^5^. Notably, preSMA (area F6) arrays were targeted to the medial wall of the right hemisphere (**Extended Data Fig. 3**), with the boundary between SMA and preSMA defined as the anterior-posterior location of the genu of the arcuate sulcus, consistent with prior studies finding significant differences across this boundary in anatomical connectivity (e.g., direct spinal projections in SMA but not preSMA^145^) and function^53,55^. In general, arrays targeted the cortical convexity immediately next to sulci, instead of within the banks, in order to allow shorter insertion depths that minimize the risk of missing the target. The exceptions were SMA and preSMA in the medial wall. To avoid damaging the superior sagittal sinus, we positioned the arrays laterally (2 mm from midline) and slanted the electrodes medially (**Extended Data Fig. 3**). All arrays were implanted in the right hemisphere (contralateral to the arm used for drawing), and are depicted in **Fig. 3** and **Extended Data Fig. 3**. The lengths of each electrode were custom designed to target half-way through the gray matter, and to vary substantially across the array, to maximize sampling of the cortical depth. Further details about electrode depths and targeted locations for the other brain regions can be found in a previous study^5^. Array connectors (Omnetics A79022) were housed in custom-made Ultem pedestals (Crist), which were secured with bone cement onto the cranial implant. Four pedestals were used per subject, holding 5, 5, 4, and 2 connectors each.

### Behavioral task

#### Task overview

The general task structure and experimental setup are described in detail in a prior publication^5^. In brief, subjects were seated comfortably in the dark with head restrained by headpost fixation. They faced a touchscreen (Elo 1590L 15” E334335, PCAP, 768 x 1024 pixels, refresh rate 60 Hz, with matte screen protector to reduce finger friction) that presented images and was drawn on (diagram in **Extended Data Fig. 1**). Both subjects decided on their own over the course of learning to perform the task with the left hand. Gravity-delivered reward (water-juice mixture) was controlled by the opening and closing of a solenoid pinch valve (Cole-Parmer, 1/8" ID). Subjects were water-regulated, with careful monitoring to ensure that consumption met the minimum requirement per day (typically exceeding it), and body weight monitored to ensure good health. The task was controlled with custom-written software, using the MonkeyLogic (2.2.45) behavioral control and data acquisition MATLAB package^146^ (PC: Windows 10 Pro, Intel Core i7-4790K, 32GB RAM; DAQ: National Instruments PCIe-6343). All stimuli (images of line figures defined as point sets, with points rendered large enough to appear as continuous curves) were also generated with custom-written MATLAB (R2021a) code.

#### Trial structure

Trial structure is summarized in **Fig. 1e** (screen schematic in **Extended Data Fig. 1**) and described in detail in a prior publication^5^, with the following timing parameters for the current study. The pre-image hold period (duration between pressing the fixation button and image onset) ranged from 0.6-0.8 s (uniformly distributed) to 1.75-2.15 s across experiments. The “plan” period (duration from image onset to go cue) ranged from 1.25-1.55 s (uniformly distributed) to 1.8-2.4 s across experiments for Monkey P and was 1.8-2.4 s for Monkey D. Screen image changes (including image presentation and other trial events) were recorded using photodiodes (Adafruit Light Sensor ALS-PT19) and sounds were recorded using an electret microphone (Adafruit Maxim MAX4466, 20Hz-20KHz). We performed eye tracking (ISCAN), but did not enforce eye fixation.

#### Grammar Task

Subjects were first trained to the point where they understood the basic task structure (to reproduce images by tracing on a touchscreen) and learned their subject-specific sets of action symbols (each mapped onto a unique shape in the current study). The procedure for this training is described in detail in a prior publication^5^.

Subjects were then trained on the Grammar Task, in general following the grammar A^n^B^m^C^k^ and in some experiments A^n^B^m^, as noted case by case in the results. The essence of the task is to reproduce an image (shape array in a grid) by tracing it with a sequence of strokes, one for each shape component, and with strokes in the correct order, which is defined by applying the rules of the learned grammar to the particular image. This task structure is described in the context of example problems in the results related to **Fig. 1**. In principle, a given image can have multiple correct sequences under the grammar, because the grammar does not specify what order to draw items within each role. To ensure that there was only one correct sequence per image, in most cases (all 16/16 recording sessions and 2/2 behavioral experiments for Monkey D and 7/13 recording sessions for Monkey P) a directional rule (e.g., draw left to right in **Fig. 1e**) was trained along with the grammar and images were designed so that no two instances of a given role had the same rank along that direction axis. For the remaining 6/13 recording sessions for Monkey P and 1/1 behavioral session, instead of imposing a rule, we allowed the monkey to express the directional bias that he expressed spontaneously, even without training. In general, subjects needed to use one stroke per image component; using more than one led to automatic and immediate failure of the trial (an “abort”). In one behavioral experiment for Monkey P we allowed the subject to use additional strokes, if desired, to “clean up” an already completed component, but the subject used one stroke per image component a large majority of the time (98% of trials). We also included a constraint to abort the trial if the drawing veered substantially off the lines of the image, in order to further incentivize consistent quality of strokes; this occurred on a small fraction of trials (mean and S.D.: 12.7% +/-6.3% trials). Because these trials were not completed, it was impossible to classify them as correct or incorrect. Therefore, these cases were excluded from behavioral analyses (below). The trajectory of the stroke was not enforced, but subjects naturally chose to use trajectories consistent with their learned action symbols^5^.

In general, in a given epoch (a contiguous duration during which a specific mapping between role and shape was unchanging), a single shape set was learned, meaning that a given role (A, B, or C) was always associated with a specific shape (e.g., A=circle, B=line, C=chevron). For analysis of role abstraction (**Fig. 4**), we trained subjects to switch between two shape sets on a trial-by-trial basis. Which shape set was active on each trial was cued by the content of the image itself.

Training on the Grammar Task involved progression through curriculum stages in a manner that was dependent on performance. There were two components to the training curriculum: decreasing stroke- by-stroke instruction (**Fig. 1f**) and increasing problem complexity (**Fig. 1g**). Stroke-by-stroke instruction refers to visual cueing of the order in which image components should be drawn. Instruction involved the following sequence of trial events (illustrated in **Fig. 1f**). First, the target image is shown. Then all components fade out to lower contrast (although still visible) except the one component that needs to be drawn next, which remains in high contrast, which we call the “guide cue.” If the subject successfully draws that component, the next component that needs to be drawn is shown in high contrast so that it is the current guide cue. This process continues until the entire image is drawn. If at any point the subject draws the incorrect component, the trial aborts (immediate white noise and change of screen color to red), and no reward is provided. For the curriculum, stroke-by-stroke instruction was decreased by progressing through stages where the components that were not the guide faded out to a decreasing extent, until at the final stage these components were not faded out at all, and thus the same contrast as the guide cue. At this point the subject had to decide on its own the correct sequence of components. We progressed manually in a performance-based manner through five stages of decreasing fade, implemented as alpha values increasing through levels of 0.05, 0.37, 0.68, 0.95, and 1.0 (no instruction).

Problem complexity is defined as the number of components in the image, summed over all shapes. This was increased over stages, where each successive stage added problems from the next level of complexity. Stages progressed starting from two components up to four with the maximum number of items set at two for A and for B (the two behavioral experiments for Monkey D) or up to six with the maximum number of items set at three for A and for B (the behavioral experiment for Monkey P).

After completing the curriculum, we tested generalization. Test problems were more complex than those seen in training and were designed to test for different possible behavioral strategies, including use of the grammar. This was done by interleaving test problems with problems from the training distribution [fraction of trials of 0.29 (Monkey D) to 0.33 (Monkey P)]. To assess generalization without “contamination” of additional learning, we needed to avoid providing feedback that may lead to further learning of a particular sequencing strategy. Therefore, for test problems, subjects were allowed to use any sequence and reward was determined by accurate completion of the image, and not the stroke sequence.

We compared monkey behavior on generalization trials to three hypotheses alternative to grammar generalization. For each hypothesis, we simulated the expected repeat length in behavior on the same problems given to the monkey. For the “Random” strategy, for each problem we sampled 100 random sequences by uniformly sampling shape components without replacement, and took the average repeat length over those sequences. For the “Direction” strategy, we sampled strokes following a spatial directional rule using the subject’s own directional bias. For experiments in which a direction rule was imposed in training (see above), this direction was used for sampling sequences. For experiments without a rule imposed in training, we used the subject’s own directional bias, quantified as the mean movement angle over all trials, where each trial’s angle was defined by first computing the mean vector over all stroke-to-stroke movements (i.e., the vector between the centroids of two shape components), then computing the mean vector over all trials, and then taking the angle of the resulting vector, and then discretizing that vector to up, down, left, or right. For images without a unique solution after applying this direction rule (e.g., if direction is left, and there are two items in a column), we took the average repeat length over all sequences after uniformly sampling sequences consistent with the direction rule. For the “Motifs” strategy, for each problem we sampled a sequence by concatenating motifs from the set of sequences experienced during training. For Monkey D, this set was {A^n^B^m^ | n<=2, m<=2} and for Monkey P this was {A^n^B^m^ | n<=3, m<=3}. A given problem’s sequence was determined through an iterative process, by taking the motif with the largest value of n and m, removing that from the image, and then repeating that process, concatenating the resulting sampled motifs into a sequence (e.g., **Fig. 2a**).

For the Single-action Task (**Fig. 3**), each image consisted of a single instance of one of the shapes involved in that day’s grammar. Single-action and Grammar Task trials were switched in blocks, and across trials, the locations of shapes in the Single-action task were varied to match the locations used for the Grammar Task.

### Behavioral data analysis

#### Preprocessing of touchscreen data

Touchscreen data were represented as time series of (x, y) coordinates in units of pixels (conversion: 33.6 pixels/cm) and sampled at 60 Hz, which we upsampled to 500 Hz (performed in MonkeyLogic to align all behavioral signals, including trial event markers and eye tracking), and low-pass filtered below the highest frequency in the drawing movements (15 Hz). Strokes were segmented based on the time of first touch (onset) and the time of last touch (offset) with 500 Hz resolution.

#### Quantifying grammar task performance

To quantify the extent to which behavior was consistent with the grammar, for a given shape we computed the repeat length in behavior and compared that value to the number of items of that shape in the image (**Fig. 1i-k, 2c**). If the grammar is being used, then these two values should be identical. If any of the three other potential strategies are used, then repeat length should be substantially lower than the number of items (**Fig. 2a**). For the behavioral experiments using the A^n^B^m^ grammar, we restricted our analysis to A. This is because producing the correct repeat lengths of B could result from a simple strategy of drawing whatever items remain not-yet-drawn in the image, because once the subject has started drawing B’s the only image components not yet drawn are B’s. Therefore, restricting analysis of repeat length to A is a stronger test of grammar use.

To quantify generalization, our analysis of repeat length ignored the direction of strokes. This is because whether a sequence is consistent with the grammar is determined by the sequence of shapes, and not by the order within each shape (e.g., the sequence AABBBC is consistent with the grammar whether it is drawn left-to-right or right-to-left). To further ensure that we were assessing generalization for test problems, we quantified performance only for the first trial each problem (unique image) was encountered.

#### Quantifying gap duration

We quantified the relationship between gap duration and sequence context, including whether this relationship was consistent with an understanding of grammar structure (**Fig. 2d-f**). For a given gap, its duration was defined as the time between the offset of its preceding stroke and the onset of its following stroke. To compare gaps across vs. within role repeats (**Fig. 2d**), we analyzed pairs of gaps, where for each pair one gap was across roles (e.g., A to B) and the other gap was within a role (e.g., A to A), and where the two gaps were measured in the same session and had the same roles (A, B or C) for their preceding strokes. We excluded gaps that skipped a role (e.g., A to C). To compare gaps that skip a role (e.g., A to C) vs. gaps between adjacent roles without skipping a role (e.g., A to B) (**Fig. 2e**), we analyzed pairs of gaps, where for each pair one gap skips a role and the other gap does not, and where the two gaps were the same gap number within the drawing sequence, and had the same roles for their preceding strokes. We included gaps that compared START to B (skips A) vs. START to A (does not skip A). To assess the effect of length of the upcoming repeat on the current gap duration (**Fig. 2f**), we compared gaps that differed in the length of the upcoming repeat, but had the same number of strokes already completed, number of strokes remaining in the trial, role of preceding stroke, and role of following stroke. Applying these constraints allowed us to isolate the effect of repeat length on gap duration, but also restricted the number of eligible cases such that we only had data for one animal (**Fig. 2f**).

We performed statistics on gap duration using multiple linear regression to estimate the effect of sequence context (the independent variable of interest) on gap duration, controlling for other variables, including distance travelled during the gap (**Extended Data Fig. 2**). Models were fit separately for each experimental session. For the analysis comparing gaps within-role vs. across-role, we modeled gap duration as a function of sequence type (indicator variable for whether the gap is within or across roles), distance travelled during gap (z-scored), role of preceding stroke, the number of preceding strokes, and the number of remaining strokes. For the analysis comparing gaps that skip a role vs. those that do not, we performed the same regression analysis, but with sequence type as an indicator variable for whether the gap skipped a role or not.

### Neural recordings

Recordings were acquired using a Tucker-Davis Technologies (TDT) system, including headstage (Z-Series 32 Channel Omnetics, LP32CH-Z), amplifier (PZ5M-256), processor (RZ2), and storage (RS4), sampled at 25 kHz (local reference mode), controlled with TDT Synapse (v98) software run on a Windows 10 PC (Intel Core i7-3770, 32GB RAM), and saved to disk. Analog and digital task-related signals, including behavioral events (photodiode, audio, and trial event markers) and eye tracking (ISCAN, 125 Hz), were synchronized to external triggers recorded by the neural data acquisition system.

### Neural data preprocessing

We extracted for later analysis both isolated single-unit (SU) and multi-unit (MU) spike clusters from the stored broadband signal following procedures using Kilosort^147^ (v2.5) for an initial spike sorting step and then applying custom-written software for curation of units. Spike times were then converted to firing rate functions by smoothing with a 0.025 s Gaussian kernel (0.01 s slide). These procedures are described in detail in a prior publication^5^.

### Neural data analyses

#### Dimensionality reduction of population activity

We performed dimensionality reduction on the neural population activity, in order to reduce the influence of noise, which can reduce the interpretability of the Euclidean distance for high-dimensional data^148^, using an approach that identifies a linear projection of population activity (i.e., a subspace) that maximally captures variance in activity due to all task-relevant variables, thereby denoising the data. The overall procedure involved “reshaping” time-varying population activity into a single vector representation, correcting activity for the gross effect of the initial reach at the start of the trial and performing dimensionality reduction to maximize variance due to task-relevant variables. The details of this procedure are as follows.

Each datapoint corresponded to population activity in a time window temporally aligned to the onset of a stroke (−0.35 s to 0.2 s relative to stroke onset), such that a session’s dataset for a given brain region was represented as a K x N x T matrix, where K is the number of strokes, N is the number of units for that region, and T is the number of time bins (0.15 s window, 0.05 s slide). To summarize this time-varying activity into a “static” representation, we concatenated the N channels’ length T vectors end-to- end to construct a matrix of size K x NT. We performed PCA on this matrix and kept the first 50 principal components, resulting for each region in a data matrix *D* of size K x 50.

We then corrected activity for the gross effect of the initial reach that initiates the drawing, moving from holding the start button towards drawing the first stroke (**Fig. 1e**). This large movement is associated with a large change in neural activity, especially in the motor and premotor areas^5^. To allow assessing encoding of grammar structure related activity, and to compare across brain areas, we applied a simple linear correction to reduce the effect of this initial reach, as in a prior publication^5^. We residualized activity to remove the mean effect on neural activity associated with being the first stroke of the trial. For each dimension of *D* (i.e., each principal component), we fit the linear regression model

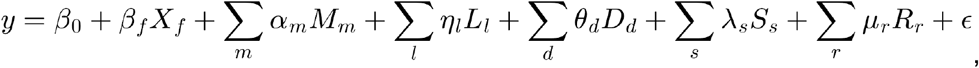

where *y* is scalar activity for a given stroke; *β*_0_ is the intercept; *X_f_* indicates whether the stroke is first (*X_f_=1*) or not (*X_f_=0*); *M_m_* are continuous kinematic covariates (location of stroke onset, displacement from hold button (or previous stroke for non-first strokes) to stroke onset, and stroke mean velocity, each in x and y); *L_l_*, *D_d_*, *S_s_* and *R_r_* are indicators for grid location (categorical), difference in grid location from the previous stroke (categorical), shape, and sequence context (a categorical variable that is the conjunction of shape, repeat length, and ordinal position within shape); and ɛ is residual error. We then subtracted from all first-stroke data the contribution of the first-stroke term, resulting in a dataset *D\** that is the same as *D* but corrected for the mean effect of being the first stroke.

We then performed dimensionality reduction to denoise the data and maximize variance due to task-relevant variables, using targeted dimensionality reduction^85,149^. For each neural dimension (column) of *D\**, we fit a linear regression model predicting activity from categorical predictors for multiple task-relevant variables. For analysis of grammar-specific roles (**Fig. 3**), the dataset combined strokes from Single-action and Grammar tasks, so we used task-relevant variables that varied across both tasks: shape and task kind. For other analyses (**Figs. 4-6**), the categorical task-relevant variables were grid location, difference in grid location, shape, and sequence context (the conjunction of shape, repeat length, and ordinal position within shape). The resulting regression coefficients across levels of all variables defined a vector of task-relevant coding. Stacking these coefficient vectors across neural dimensions yielded a coefficient matrix of size N x L, where N is the number of neural dimensions retained after the initial PCA (50), and L is the total number of coefficients contributed across variables (each variable contributed a number of coefficients equal to the number of levels for that variable minus 1). Conceptually, this matrix can be understood as the directions in the space of population activity along which each task variable is encoded. For analysis of grammar-specific roles (**Fig. 3**), we wished to identify a shape encoding subspace that is invariant to task kind, if it exists, so we kept only the columns of that matrix corresponding to shape coefficients. For the other analyses, we kept all columns. To identify the final task-variable encoding subspace, we performed PCA on this coefficient matrix and retained the top 6 principal components, orthonormalized using QR decomposition, resulting in a basis set defining the task-variable encoding subspace. Population activity (*D\**), projected onto this subspace, was used as input data for subsequent population state space plots and analyses of neural distance.

#### Computing the neural distance metric

To summarize the encoding strength of select variables in population activity, including of shape (**Fig. 3, 4**) and ordinal position (**Fig. 5**), we computed the neural distance metric, as defined in our prior study^5^. It quantifies the dissimilarity of population activity between two sets of strokes of condition A and B, where each condition is a specific conjunctive value of task-relevant variables, matched in all variables except the one variable whose effect on neural activity is being tested. In brief, inspired by the “normalized distance” in Liu et al^150^, it is the average pairwise Euclidean distance across strokes in conditions A and B, minus the average within-condition distance. This subtraction ensures the useful property that this distance is unbiased, in that the expected value of neural distance between two sets of strokes sampled from the same distribution is zero (unlike the mean Euclidean distance, which is biased upwards^151^). The resulting distance is divided by an upper-bound to normalize it between 0 and 1. Neural distance is defined as:

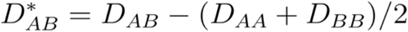

where the normalized Euclidean distance between sets of strokes (indexed by *i* and *j*) in conditions *A* and *B* is:

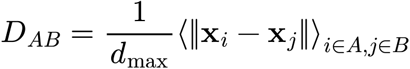

Here, x is the population activity vector after applying targeted PCA, and *d_max_* is an upper-bound (98th percentile) of the distances between all pairs of different strokes combined across all conditions, and ||.|| is the 2-norm.

#### Quantifying shape separation

To quantify how strongly shape was encoded in population activity (“shape separation” in **Figs. 3, 4**), we computed the mean effect of shape on population activity, using the neural distance metric, while controlling for other task-relevant variables. Shape separation was the average pairwise neural distance between stroke conditions differing by shape, computed for specific subsets of the data as appropriate for each analysis. For **Fig. 3h-j**, shape separation was computed for just the Single-action task (y-axis) or just the Grammar task (x-axis). For **Fig. 4h-j**, we either used stroke pairs that had the same role, ordinal position, and repeat length, but were from different shape sets (y-axis), or stroke pairs that were from the same shape set (and had the same repeat length and either the same or different ordinal position) but had different roles (x-axis). In all the above cases, we controlled for other behavioral covariates that may affect neural activity by using a stratified analysis. The strokes were partitioned into strata defined by unique combinations of the behavioral covariates, shape separation was computed independently within each stratum (as described above) and then averaged over strata. The behavioral covariates were the categorical variables of grid location and grid location of the previous stroke (which controls for the movement vector during the gap).

To compare brain regions in their relative encoding of shape separation in the Grammar task (x-axis, **Fig. 3h**) vs. the Single-action task (y-axis, **Fig. 3h**), we used a linear regression based method to model the overall relationship between these two metrics across brain areas, and then computed each area’s deviation from that model as a quantification of role encoding. Within each session, we fit an ordinary least-squares line relating y to x across brain-region means *y=a+bx*. For each region with coordinates *(x_0_, y_0_)*, we then computed the signed orthogonal residual from the fitted line

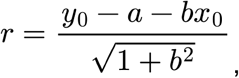

with negative r indicating that the region lies below the best-fit line. We also computed the signed projection along the line.

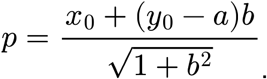

This procedure was performed in the same manner to test for role abstraction (**Fig. 4**).

#### Quantifying ordinal separation

Computed analogously to shape separation, ordinal separation (**Fig. 5f, g**) was the average pairwise neural distance between sets of strokes that differed in ordinal position, and that matched in role and repeat length. We controlled for the behavioral covariates of grid location, and grid location of preceding stroke using a stratified analysis as described for shape separation.

To assess whether ordinal encoding was more consistent with “counting up” versus “counting down” (**Fig. 5h-j**), we performed the following analysis. Conditioning on each unique sequence context (x-axis in **Fig. 5i**), defined by the combination of role, repeat length, ordinal position relative to repeat start (ord_FWD_; grouped into 1, 2, and 3 or higher), and ordinal position relative to repeat end (ord_REV_; grouped into −3 and lower, −2, and −1), we computed two neural distances, one to all other strokes of the same role that have different ord_REV_ and the same ord_FWD_ (orange curve in **Fig. 5i**), and the other to all other strokes of the same role that have different ord_FWD_ and the same ord_REV_ (blue curve in **Fig. 5i**). In both cases, neural distance was computed only between stroke pairs with matching grid location, and grid location of previous stroke.

To test whether gap duration relates to the extent to which ordinal encoding was more strongly “counting up” vs. “counting down” (**Fig. 5k**) we did the following. In a given session, mean gap duration was computed for each role by first averaging over all trials for each ordinal position and then over all ordinal positions. We computed two neural distances for each role. One was the distance between pairs of strokes with different ord_FWD_ but same ord_REV_, and the other was for pairs with different ord_REV_ but same ord_FWD_, in both cases computed using a stratified analysis with behavioral covariates of grid location and previous grid location and averaged over all ordinal positions. We then took the difference of these two neural distances to derive the relative encoding of ord_FWD_ and ord_REV_ (x-axis in **Fig. 5k**).

#### Computing angles between variable-encoding axes

We quantified the geometry of population activity (**Fig. 6**) in the following manner. We used data after dimensionality reduction (above), using the top six principal components from targeted PCA. We then assessed geometry based on quantifying the pairwise angles for select pairs of condition-difference vectors. Each vector was computed as difference of trial-mean activity between two conditions, where each condition was a unique combination of the following task-relevant variables: role, repeat length, ordinal position, grid location, preceding grid location, such that a vector between the conditions (*c*_1_, *c*_2_) was

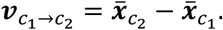

Where **x̄**_c_ is average activity over strokes. We excluded conditions with fewer than four datapoints (strokes). We computed two kinds of vectors:

1. Within-role ordinal vectors. The two conditions that define this vector differed in ordinal position and matched in all of the other task-relevant variables. One mean vector was computed for each combination of those other task-relevant variables (i.e., each grouping variable). We retained only condition pairs in which ordinal position increased between *c*_1_ and *c*_2_, and only adjacent levels of ordinal position after sorting the observed levels. Collecting the resulting vectors for each pair of ordinal positions for each role (e.g., 1◊2) yielded one set of vectors for that pair (differing in the grouping variable).
2. Across-role vectors. Condition pairs differed in role and were matched in all of the other task-relevant variables. One mean vector was computed for each combination of those other task-relevant variables. We retained only condition pairs in which role increased between *c*_1_ and *c*_2_, and only adjacent levels of role after sorting the observed levels (e.g., A, B, C). Collecting the resulting vectors for each pair of roles yielded one set of vectors (differing in the grouping variable).

Given these two sets of vectors, we then defined three types of vector pairs:

1. Ordinal. Both vectors in this pair type were within-role ordinal vectors that were in the same role.

For example, if the set of within-role ordinal vectors were for the ordinal pairs {1◊2, 2◊3, 3◊4}, then the ordinal pairs were {(1◊2, 2◊3), (1◊2, 3◊4), (2◊3, 3◊4)}.

1. Role. Both vectors were across-role vectors. For example, if the across-role vectors were {A◊B, B◊C}, then the role pair was (A◊B, B◊C).
2. Ordinal (different role). Both vectors were within-role ordinal vectors that were in different roles.

For each of the three vector pair types, we then computed the mean pairwise vector angle in the following manner. Each vector pair type consisted of a set {(A, ℬ)} where each (A, ℬ) is a specific pair of sets of vectors, with 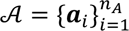 and 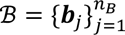 (e.g., all 1 → 2 within-role ordinal vectors differing by grouping variable and all 2 → 3 within-role ordinal vectors differing by grouping variable). We computed the across-set and within-set mean dot products

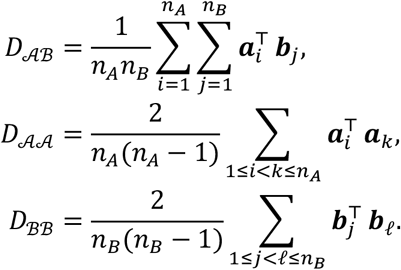

The self-products exclude 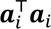 and 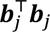, and therefore provide cross-validated estimates of squared norms. Cases lacking at least two vectors in either set (so that *D*_AA_ or *D*_ℬℬ_ is undefined) were excluded, such that each (A, ℬ) yielded one triple (*D*_Aℬ_, *D*_AA_, *D*_ℬℬ_) for each grouping level.

Means over *D*_Aℬ_, *D*_AA_, and *D*_ℬℬ_ were taken by first, within each session, averaging over all grouping levels, and then averaging over all experiments to get a single set of scalar values for each vector pair type for *D*_Aℬ_, *D*_AA_, and *D*_ℬℬ_. We then computed cosine similarity from the three mean dot products,

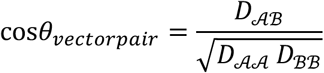

To be able to assess vector pair types involving across-role vectors (the pair type “role”), we analyzed only experiments with at least three roles. This included all experiments for monkey D and a subset of monkey P. For analysis of the vector pair type “ordinal (different role)”, we also included cases with two roles (only if data for at least two ordinal positions with sufficient data existed per role). We performed the following statistical tests of the hypothesis of high ordinal cosine similarity compared to the other two vector pair types, by directly contrasting ordinal vs. each of the other vector pairs,

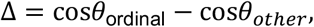

where other was either “role” or “ordinal (different role)”. Significance of Δ was assessed with a two-sided permutation test (*N* = 10,000). Within each animal, date, and brain region, labels of low-level vector-pair datapoints—each datapoint an intact triple (*D*_Aℬ_, *D*_AA_, *D*_ℬℬ_) for a given grouping level—were randomly remapped across each other, preserving the triple, but shuffling the triple’s label in terms of its vector pair type. The test statistic was recomputed after each remapping, and the p-value was taken as the two-sided fraction of null values at least as extreme as the observed Δ.

## Data and code availability

The data and code used in this study are available upon reasonable request.

## Acknowledgements

We thank A. Rouse, M. Eldridge, and M. Schieber for assistance with surgeries; K. Panikhina, X. Ma, and other members of the Freiwald, Wang, and Tenenbaum labs for project feedback; T. Nigam, J. Rhee, C. Correa, Y. Prut, and K. Kay for manuscript feedback; V. Sherman and A. Gonzalez for technical assistance; and L. Yin for administrative assistance. This work was supported by the National Institutes of Health, through the National Eye Institute (R01EY021594 to W.A.F.), the National Institute Of Mental Health (F32MH125573 to L.Y.T.), and the National Institute Of Neurological Disorders And Stroke (K99NS131585 to L.Y.T.), as well as the Simons Foundation’s Collaboration on the Global Brain (876120SPI and AN-NC-GB-Pilot Extension-00002596-01 to X.-J.W., J.B.T., and W.A.F., and NC-GB-CULM-00003138 to X.-J.W.), the Center for Brains, Minds & Machines of the National Science Foundation (STC award CCF-1231216 to W.A.F. and J.B.T.), the Office of Naval Research (N00014-20-1-2292, Vannevar Bush Faculty Fellowship, to W.A.F., N00014-23-1-2040 to X.-J.W., and MURI N00014-21-1-2801 to J.B.T.), the Air Force Office of Scientific Research (FA9550-22-1-0387 to J.B.T.), and the Price Family Center for the Social Brain (W.A.F.).

## Author contributions

Study conception and design: L.Y.T., X.-J.W., J.B.T., and W.A.F. Animal task training: L.Y.T., with input from D.J.H. and K.G.G. Experiments: L.Y.T., D.J.H., and K.G.G. Data analysis: L.Y.T., with input from D.J.H., K.G.G., and Y.L. Manuscript initial draft: L.Y.T. Manuscript revision: All authors. Funding acquisition: L.Y.T., X.-J.W., J.B.T., and W.A.F. Project supervision: X.-J.W., J.B.T., and W.A.F.

---

**Extended Data Fig. 1.**
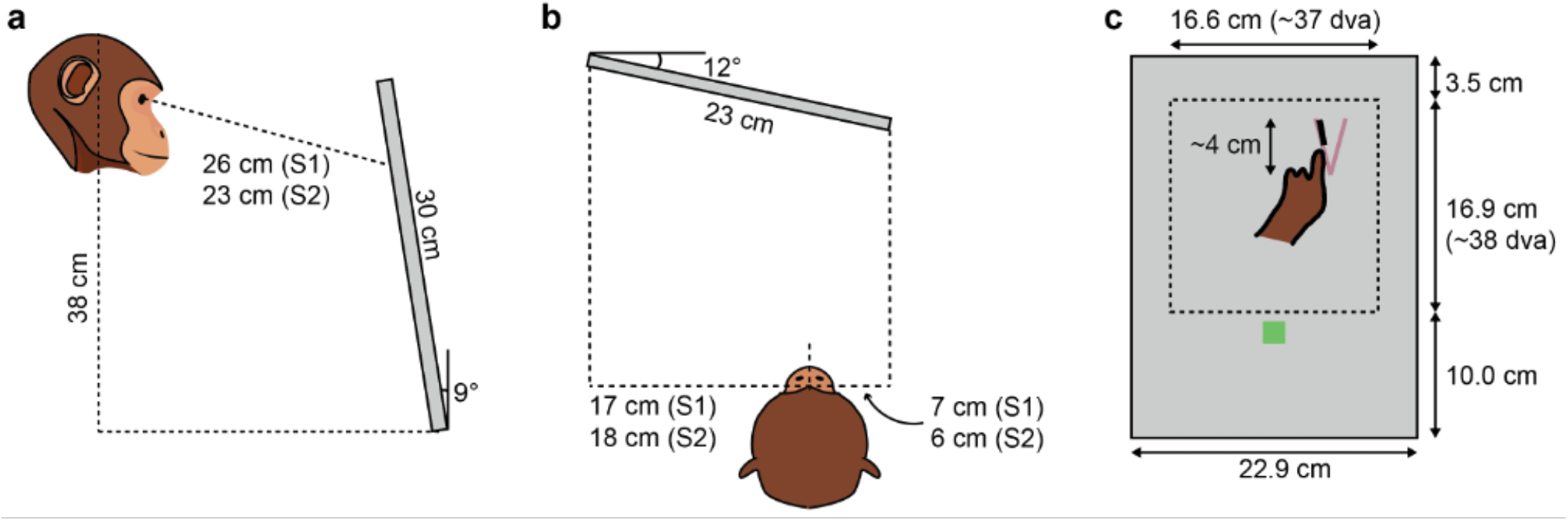
Behavioral task setup. (a) Schematic of subject relative to touchscreen, profile view, during recording sessions. Monkeys P (S1) and D (S2) were positioned at different distances to accommodate their individual anatomies and postures. The screen was slanted slightly to optimize the ability to see and reach to the same part of the screen (the workspace at the top of the screen; see panel c). (b) Schematic of subject relative to screen, top view. Subjects were positioned to the right to accommodate reaching with the hand used for drawing (left). (c) Schematic of screen during trial, with component locations and sizes to scale. The finger is tracing over the figure (purple-gray “V”), leaving a black trail of “ink” behind. The average size of shapes was 4.0 cm (maximum of width and height). The thickness and color of the figure and “ink” are to scale. The subject can press the “done” button (green square) at any time to report completion. Dashed line (not visible to the subject) indicates the workspace where images and drawings occurred.

**Extended Data Fig. 2.**
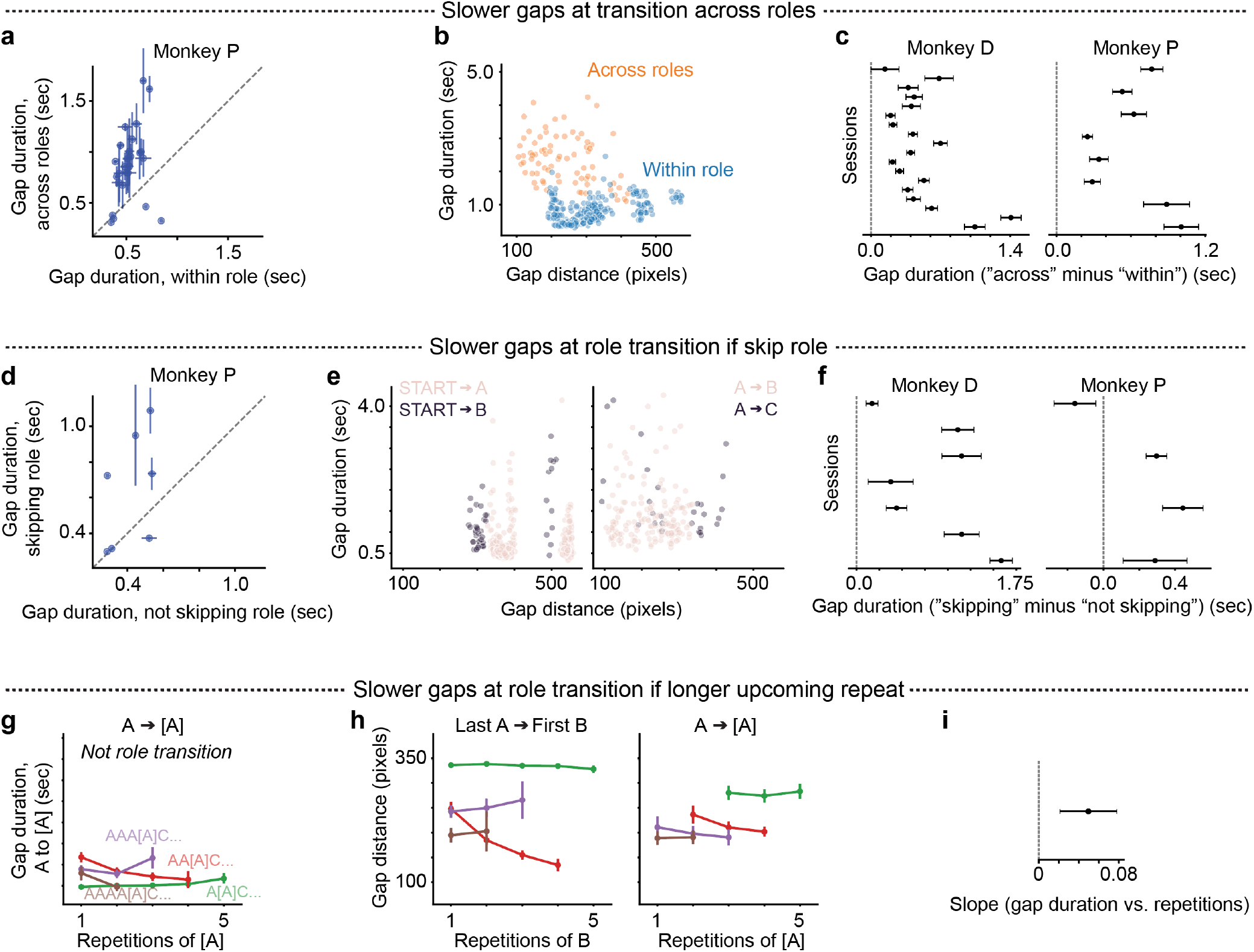
Duration of inter-stroke gaps reflects grammatical structure. (a) Gap duration reflects the hierarchical structure of the grammar. Quantification of mean gap duration for transitions across roles (y-axis) vs. within roles (x-axis). Same as Fig. 2d, but for monkey P. (b) Gap duration reflects the hierarchical structure of the grammar, even after controlling for gap distance. Gap duration across (orange) and within (blue) roles, as a function of the distance traveled (displacement from end of previous stroke to start of next stroke). Shown are data for an example case (AB vs. AA, gap between strokes 1 and 2) for monkey D. (c) Summary of effect on gap duration (across minus within roles) controlling for other factors that may influence gap duration. This shows regression coefficients (with 95% CI) for “across” minus “within”, using multiple linear regression predicting gap duration with regressors of gap distance, role preceding gap, and number of upcoming strokes (Methods). (d) Gap duration reflects the order of roles (A to B to C). Quantification of gap duration for transitions skipping a role (y-axis) vs. not skipping a role (x-axis). Same as Fig. 2e, but for monkey P. This shows cases where the y-value is START-B and x-value is START-A. (e) Gap duration reflects understanding of the order of roles (A to B to C), even after controlling for gap distance. This shows gap duration skipping a role (dark grey) and not skipping a role (pink), as a function of the distance traveled, for two cases on the same day for monkey D, either for transitions from START (left) or from A (right). (f) Summary of effect on gap duration (skipping minus not skipping role), controlling for other factors that may influence gap duration. This shows regression coefficients (with 95% CI) for “skipping” minus “not skipping”, using multiple linear regression predicting gap duration with regressors of gap distance, role preceding gap, and number of upcoming strokes (Methods). (g) A control for the analysis showing that gap duration increases with longer repetition of the upcoming role, B (Fig. 2f); this plot shows that gap duration does not increase with longer repetition of A. These are sequence contexts matching those in Fig. 2f except that the gap duration is measured for a transition from A to A (AA) rather than to B (AB). (h) Gap distance does not increase with increasing repetitions of B (left) or A (right). This plot shows that the effect we see of increasing gap duration with increasing repetitions of B (Fig. 2f) is not an artifact of increasing gap distance. This shows the same sequential contexts as in Fig. 2f. (i) Summary of effect on gap duration of number of repetitions. This plots the regression coefficient (with 95% CI) for the effect of number of repetitions, using multiple linear regression predicting gap duration with regressors of gap distance, role preceding gap, role following gap, number of preceding strokes, and number of upcoming strokes (Methods).

**Extended Data Fig. 3.**
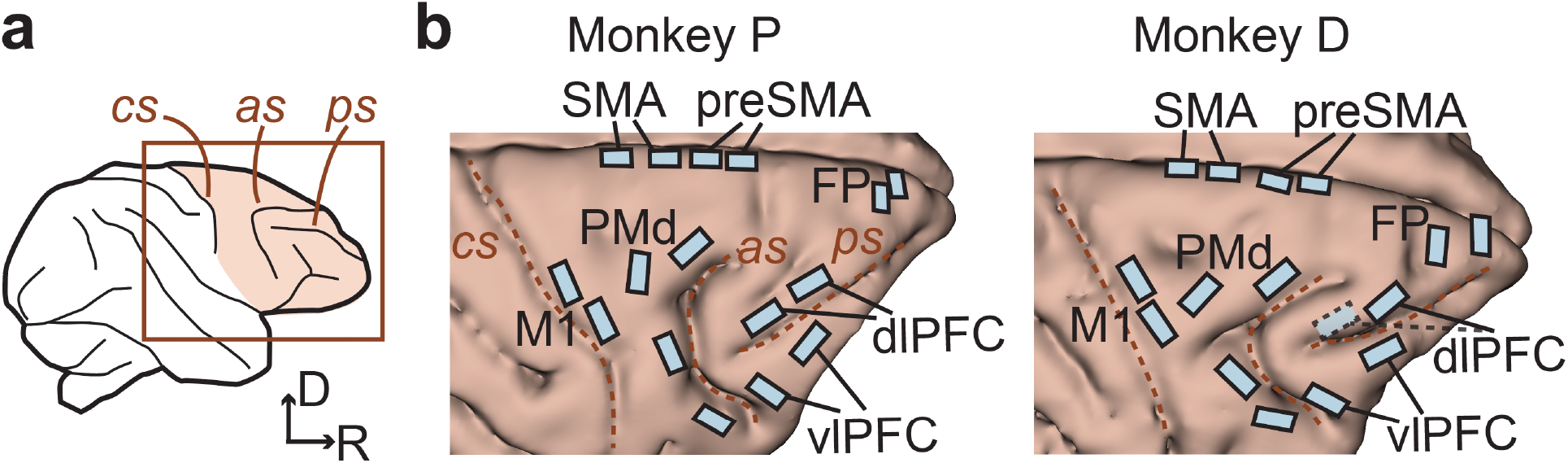
Multi-area neuronal recordings across frontal cortex and angled implantation of medial wall arrays. (a) Recordings were targeted to right frontal cortex, contralateral to the drawing hand. cs, central sulcus; as, arcuate sulcus; ps, principal sulcus. R, rostral. D, dorsal. (b) Arrays to scale on 3D rendering of brain. The caudal dlPFC array for monkey D malfunctioned. SMA and preSMA electrodes were angled to target the medial wall (see Fig. 3b). Panel b (monkey P) is same as Fig. 3a.

**Extended Data Fig. 4.**
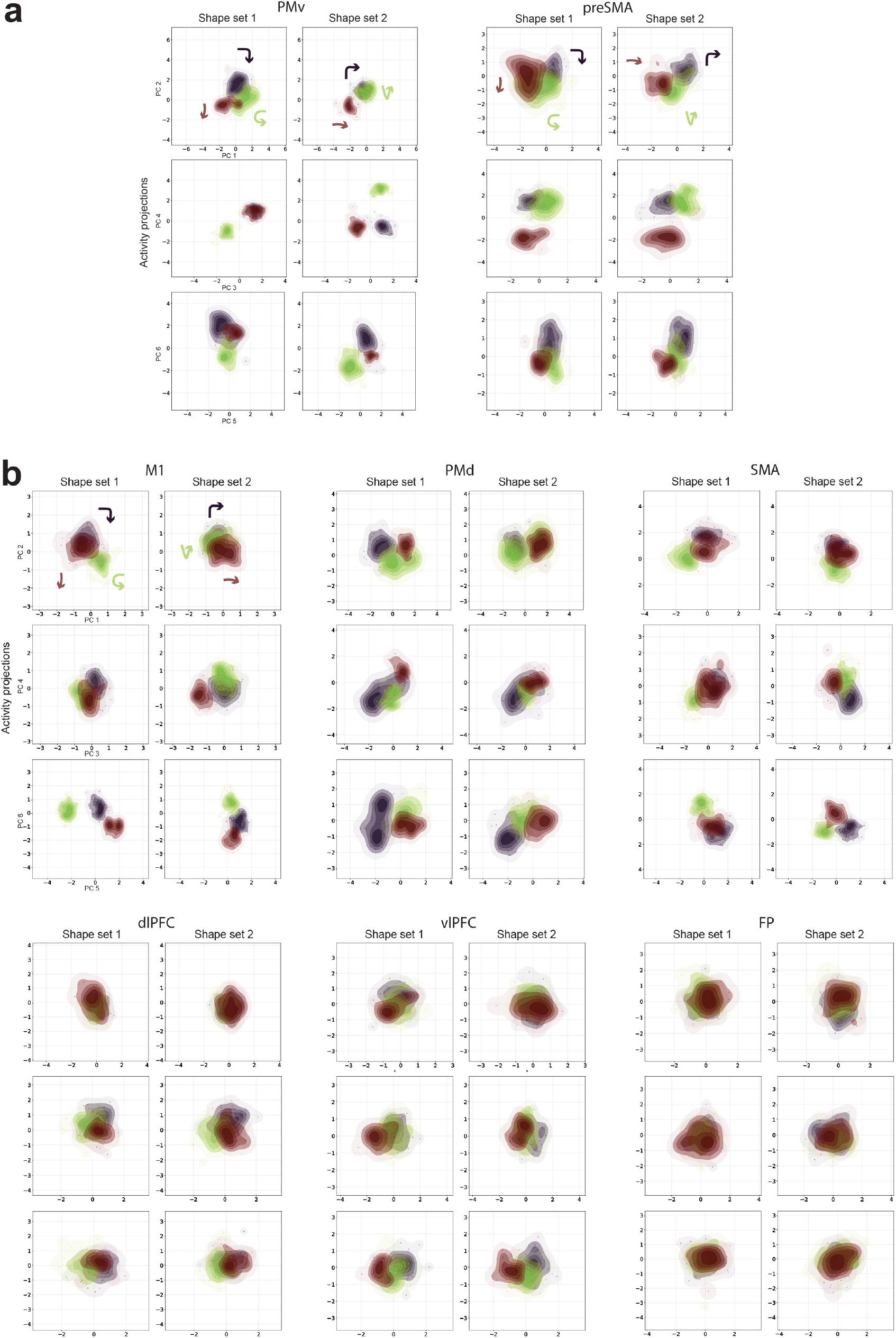
Population activity across all eight regions assessing role abstraction over specific shapes (Related to. Fig. 4). (a) Population activity for PMv and preSMA, projected to three 2D subspaces (rows) spanning the first six principal components, split by shape set (columns). Each plot shows a KDE and individual datapoints for each shape (color) in a given shape set, using data in the time window −0.35 to 0.2 sec relative to stroke onset. For each region, data are combined between shape sets before PCA and then data from both shape sets are projected into the same subspaces. The first four PCs for PMv and preSMA are also plotted in Fig. 4f, g. (b) Population activity for the other six simultaneously recorded brain regions, presented in the same manner as panel a.

**Extended Data Fig. 5.**
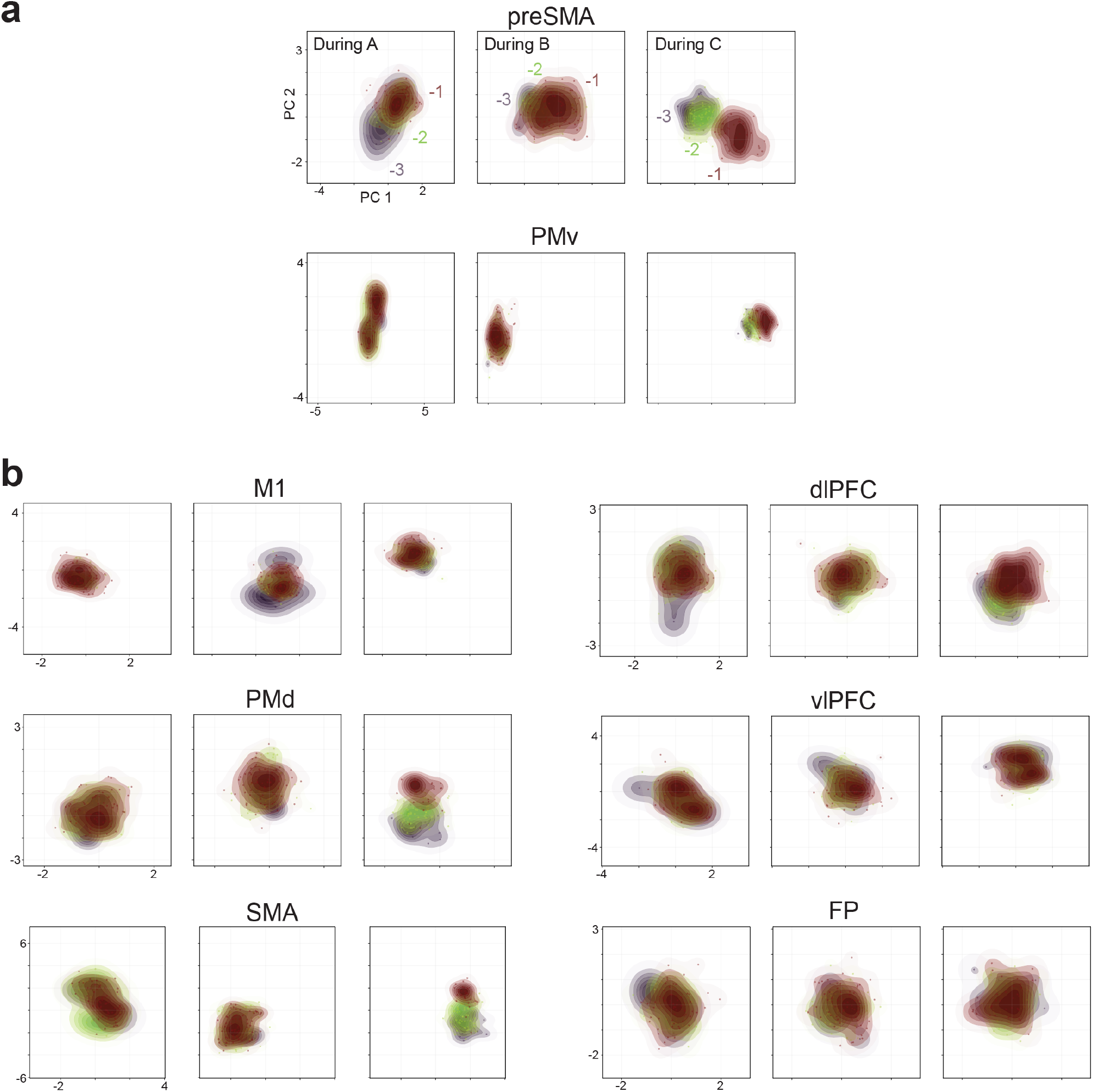
Encoding of repetition ordinal position (related to. Fig. 5). (a) Population activity for preSMA (top) and PMv (bottom) colored by ordinal position during repeats of roles A (left), B (middle), or C (right), each region plotted in the same subspace (first two principal components using PCA) across roles. Shown are only data for strokes drawn at a single specific screen location, in order to control for movement-related confounds associated with location. (b) Same as panel a, but for the six other brain regions recorded in the same experimental session.

**Extended Data Fig. 6.**
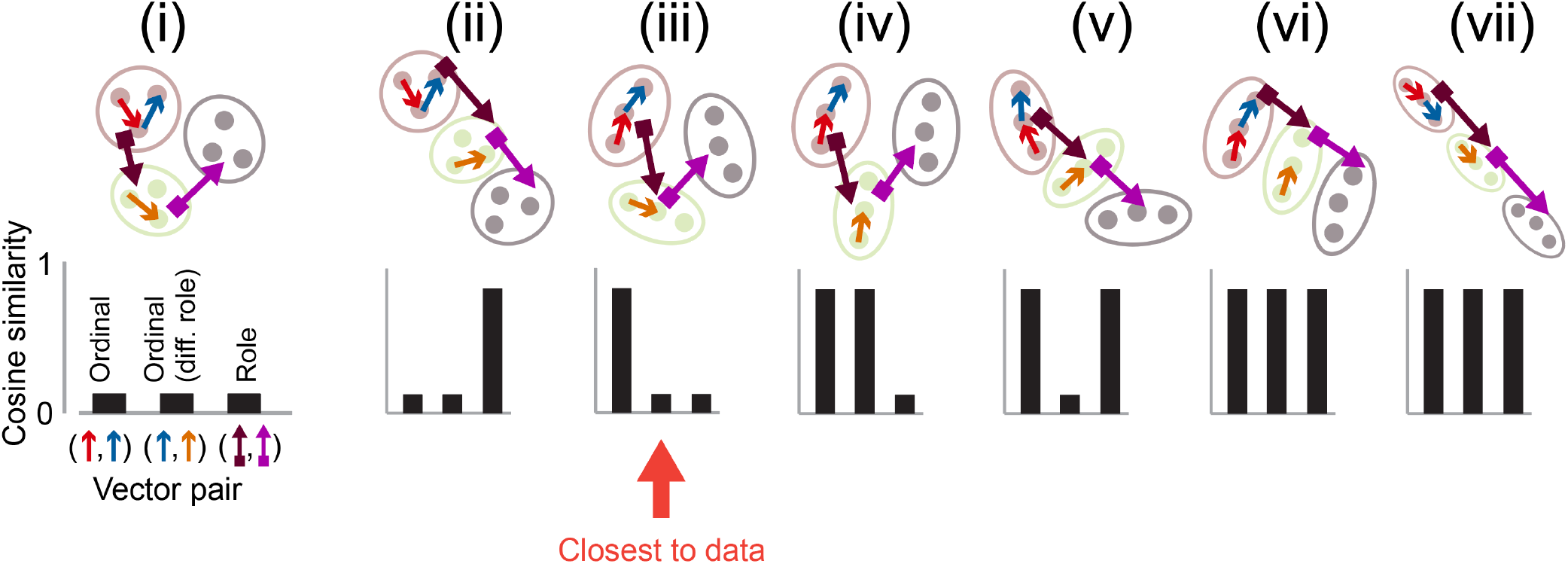
Schematic of seven potential relational structures in the geometry of population activity (related to. Fig. 6). Schematic of seven candidate population geometries (i-vii, top), with roles (A, B, C) depicted in three colors and ordinal positions (1, 2, 3) depicted as small, filled circles (see Fig. 6a for legend). Each arrow represents a vector between two states. Bottom: Predicted cosine similarity for the three kinds of vector pairs (same as in Fig. 6). These geometries differ in three major ways. First, ordinal positions may be organized along linear axes (iii-vii). This predicts high alignment between vectors taken between successive ordinal positions (high cosine similarity for “Ordinal” vector pairs). Alternatively, ordinal positions may be unstructured (i, ii), predicting low “Ordinal” vector alignment. Second, roles may be organized along a linear axis (ii, v, vi, vii; predicting high alignment for “Role” vector pairs) or unstructured (i, iii, iv). Third, linear ordinal axes, if they exist, may be parallel across roles (iv, vi, vii; predicting high alignment for “Ordinal (different role)” vector pairs) or orthogonal (iii, v). At one extreme, both ordinal positions and roles may be unstructured (i); at another extreme, they both fall along a single linear axis, consistent with a single “global” ordinal position axis (vii). We found that preSMA activity was most similar to the third structure [compare (iii) to Fig. 6].

## References

1. Kohler, W. The Mentality of Apes. (Trans E. Winter). vi, 336 (Liveright, Oxford, England, 1976).

2. Lake, B. M., Ullman, T. D., Tenenbaum, J. B. & Gershman, S. J. Building machines that learn and think like people. Behav. Brain Sci. 40, e253 (2017).

3. Boden, M. A. The Creative Mind: Myths and Mechanisms. (Routledge, 2004).

4. Ferrigno, S. Sequencing, Artificial Grammar, and Recursion in Primates. in Primate Cognitive Studies (eds Schwartz, B. L. & Beran, M. J.) 260–290 (Cambridge University Press, Cambridge, 2022). doi:10.1017/9781108955836.011.

5. Tian, L. Y. et al. Neural representation of action symbols in primate frontal cortex. Nature 1–11 (2026) doi:10.1038/s41586-026-10297-x.

6. Russin, J., McGrath, S. W., Williams, D. J. & Elber-Dorozko, L. From Frege to chatGPT: Compositionality in language, cognition, and deep neural networks. Preprint at 10.48550/arXiv.2405.15164 (2024).

7. Frankland, S. M. & Greene, J. D. Concepts and Compositionality: In Search of the Brain’s Language of Thought. Annu. Rev. Psychol. 71, 273–303 (2020).

8. Riveland, R., Pouget, A. & Driscoll, L. The compositionality continuum as a principle for studying the neural basis of intelligence. Nat. Neurosci. 1–14 (2026) doi:10.1038/s41593-026-02382-1.

9. Yang, G. R., Joglekar, M. R., Song, H. F., Newsome, W. T. & Wang, X.-J. Task representations in neural networks trained to perform many cognitive tasks. Nat. Neurosci. 22, 297–306 (2019).

10. Dehaene, S., Meyniel, F., Wacongne, C., Wang, L. & Pallier, C. The Neural Representation of Sequences: From Transition Probabilities to Algebraic Patterns and Linguistic Trees. Neuron 88, 2–19 (2015).

11. Dehaene, S., Al Roumi, F., Lakretz, Y., Planton, S. & Sablé-Meyer, M. Symbols and mental programs: a hypothesis about human singularity. Trends Cogn. Sci. 26, 751–766 (2022).

12. Quilty-Dunn, J., Porot, N. & Mandelbaum, E. The best game in town: The reemergence of the language-of-thought hypothesis across the cognitive sciences. Behav. Brain Sci. 46, e261 (2023).

13. Marcus, G. F. The Algebraic Mind: Integrating Connectionism and Cognitive Science. (MIT Press, 2003).

14. Ellis, K. et al. DreamCoder: growing generalizable, interpretable knowledge with wake–sleep Bayesian program learning. Philos. Trans. R. Soc. Math. Phys. Eng. Sci. 381, 20220050 (2023).

15. Dehaene, S., Sablé-Meyer, M. & Ciccione, L. Origins of numbers: a shared language-of-thought for arithmetic and geometry? Trends Cogn. Sci. 29, 526–540 (2025).

16. Tian, L., Ellis, K., Kryven, M. & Tenenbaum, J. Learning abstract structure for drawing by efficient motor program induction. in Advances in Neural Information Processing Systems vol. 33 2686– 2697 (Curran Associates, Inc., 2020).

17. Hayashi, M. Perspectives on object manipulation and action grammar for percussive actions in primates. Philos. Trans. R. Soc. B Biol. Sci. 370, 20140350 (2015).

18. Lake, B. M., Salakhutdinov, R. & Tenenbaum, J. B. Human-level concept learning through probabilistic program induction. Science 350, 1332–1338 (2015).

19. Karmiloff-Smith, A. Constraints on representational change: Evidence from children’s drawing. Cognition 34, 57–83 (1990).

20. Zuidema, W. & De Boer, B. The evolution of combinatorial structure in language. Curr. Opin. Behav. Sci. 21, 138–144 (2018).

21. Kazanina, N. & Poeppel, D. The neural ingredients for a language of thought are available. Trends Cogn. Sci. 27, 996–1007 (2023).

22. Gallistel, C. R. Prelinguistic Thought. Lang. Learn. Dev. 7, 253–262 (2011).

23. Cheney, D. L. & Seyfarth, R. M. Baboon Metaphysics: The Evolution of a Social Mind. (University of Chicago Press, Chicago, IL, 2008).

24. Herman, L. M., Uyeyama, R. K. & Pack, A. A. Bottlenose dolphins understand relationships between concepts. Behav. Brain Sci. 31, 139–140 (2008).

25. Stout, D., Chaminade, T., Apel, J., Shafti, A. & Faisal, A. A. The measurement, evolution, and neural representation of action grammars of human behavior. Sci. Rep. 11, 13720 (2021).

26. Rule, J. S., Tenenbaum, J. B. & Piantadosi, S. T. The Child as Hacker. Trends Cogn. Sci. 24, 900– 915 (2020).

27. Nieder, A. Prefrontal cortex and the evolution of symbolic reference. Curr. Opin. Neurobiol. 19, 99– 108 (2009).

28. Newell, A. & Simon, H. A. Computer Science as Empirical Inquiry: Symbols and Search. in (ed. Haugeland, J.) 81–110 (ACM, 1976). doi:10.7551/mitpress/4626.003.0004.

29. Goodnow, J. J. & Levine, R. A. “The grammar of action”: Sequence and syntax in children’s copying. Cognit. Psychol. 4, 82–98 (1973).

30. Byrne, R. W. & Russon, A. E. Learning by imitation: A hierarchical approach. Behav. Brain Sci. 21, 667–684 (1998).

31. Bahle, A. H. & Long, M. A. Language-like properties in the vocal behavior of animals. Curr. Opin. Neurobiol. 99, 103208 (2026).

32. Liao, D. A., Brecht, K. F., Johnston, M. & Nieder, A. Recursive sequence generation in crows. Sci. Adv. 8, eabq3356 (2022).

33. Jiang, X. et al. Production of Supra-regular Spatial Sequences by Macaque Monkeys. Curr. Biol. 28, 1851–1859.e4 (2018).

34. Whittington, J. C. R., Dorrell, W., Behrens, T. E. J., Ganguli, S. & El-Gaby, M. A tale of two algorithms: Structured slots explain prefrontal sequence memory and are unified with hippocampal cognitive maps. Neuron 113, 321–333.e6 (2025).

35. Hummel, J. E. Getting symbols out of a neural architecture. Connect. Sci. 23, 109–118 (2011).

36. Smolensky, P. Tensor product variable binding and the representation of symbolic structures in connectionist systems. Artif. Intell. 46, 159–216 (1990).

37. McCoy, T., Tenenbaum, A. L., Enyan, Z. & Zhou, Z. H. Reconciling symbols and neural networks: evidence for a quasi-symbolic view. Curr. Opin. Behav. Sci. 72, 101686 (2026).

38. Lake, B. M. & Baroni, M. Human-like systematic generalization through a meta-learning neural network. Nature 623, 115–121 (2023).

39. Tervo, D. G. R., Tenenbaum, J. B. & Gershman, S. J. Toward the neural implementation of structure learning. Curr. Opin. Neurobiol. 37, 99–105 (2016).

40. Piantadosi, S. T. et al. Why concepts are (probably) vectors. Trends Cogn. Sci. 28, 844–856 (2024).

41. Kriete, T., Noelle, D. C., Cohen, J. D. & O’Reilly, R. C. Indirection and symbol-like processing in the prefrontal cortex and basal ganglia. Proc. Natl. Acad. Sci. 110, 16390–16395 (2013).

42. Papadimitriou, C. H., Vempala, S. S., Mitropolsky, D., Collins, M. & Maass, W. Brain computation by assemblies of neurons. Proc. Natl. Acad. Sci. 117, 14464–14472 (2020).

43. Griffiths, T. L., Lake, B. M., McCoy, R. T., Pavlick, E. & Webb, T. W. Whither symbols in the era of advanced neural networks? Preprint at 10.48550/arXiv.2508.05776 (2025).

44. Binz, M. et al. A foundation model to predict and capture human cognition. Nature https://doi.org/10.1038/s41586-025-09215-4 (2025) doi:10.1038/s41586-025-09215-4.

45. Wang, L., Uhrig, L., Jarraya, B. & Dehaene, S. Representation of Numerical and Sequential Patterns in Macaque and Human Brains. Curr. Biol. 25, 1966–1974 (2015).

46. Wilson, B. et al. Auditory sequence processing reveals evolutionarily conserved regions of frontal cortex in macaques and humans. Nat. Commun. 6, 8901 (2015).

47. Zimnik, A. J. & Churchland, M. M. Independent generation of sequence elements by motor cortex. Nat. Neurosci. 24, 412–424 (2021).

48. Mizes, K. G. C., Lindsey, J., Escola, G. S. & Ölveczky, B. P. Dissociating the contributions of sensorimotor striatum to automatic and visually guided motor sequences. Nat. Neurosci. 26, 1791–1804 (2023).

49. Tian, Z. et al. Mental programming of spatial sequences in working memory in the macaque frontal cortex. Science 385, eadp6091 (2024).

50. Ohbayashi, M., Ohki, K. & Miyashita, Y. Conversion of Working Memory to Motor Sequence in the Monkey Premotor Cortex. Science 301, 233–236 (2003).

51. Barone, P. & Joseph, J.-P. Prefrontal cortex and spatial sequencing in macaque monkey. Exp. Brain Res. 78, (1989).

52. Ninokura, Y., Mushiake, H. & Tanji, J. Integration of Temporal Order and Object Information in the Monkey Lateral Prefrontal Cortex. J. Neurophysiol. 91, 555–560 (2004).

53. Nakamura, K., Sakai, K. & Hikosaka, O. Effects of Local Inactivation of Monkey Medial Frontal Cortex in Learning of Sequential Procedures. J. Neurophysiol. 82, 1063–1068 (1999).

54. Shima, K., Isoda, M., Mushiake, H. & Tanji, J. Categorization of behavioural sequences in the prefrontal cortex. Nature 445, 315–318 (2007).

55. Tanji, J. Sequential Organization of Multiple Movements: Involvement of Cortical Motor Areas. Annu. Rev. Neurosci. 24, 631–651 (2001).

56. Averbeck, B. B., Chafee, M. V., Crowe, D. A. & Georgopoulos, A. P. Neural activity in prefrontal cortex during copying geometrical shapes: I. Single cells encode shape, sequence, and metric parameters. Exp. Brain Res. 150, 127–141 (2003).

57. Averbeck, B. B., Sohn, J.-W. & Lee, D. Activity in prefrontal cortex during dynamic selection of action sequences. Nat. Neurosci. 9, 276–282 (2006).

58. Mushiake, H., Saito, N., Sakamoto, K., Itoyama, Y. & Tanji, J. Activity in the Lateral Prefrontal Cortex Reflects Multiple Steps of Future Events in Action Plans. Neuron 50, 631–641 (2006).

59. Berdyyeva, T. K. & Olson, C. R. Rank Signals in Four Areas of Macaque Frontal Cortex During Selection of Actions and Objects in Serial Order. J. Neurophysiol. 104, 141–159 (2010).

60. Manakov, M. et al. Universal Geometry of Compositional Construction in Prefrontal Cortex. 2026.04.23.720375 Preprint at 10.64898/2026.04.23.720375 (2026).

61. Sawamura, H., Shima, K. & Tanji, J. Numerical representation for action in the parietal cortex of the monkey. Nature 415, 918–922 (2002).

62. Geddes, C. E., Li, H. & Jin, X. Optogenetic Editing Reveals the Hierarchical Organization of Learned Action Sequences. Cell 174, 32–43.e15 (2018).

63. Ferrigno, S., Cheyette, S. J., Piantadosi, S. T. & Cantlon, J. F. Recursive sequence generation in monkeys, children, U.S. adults, and native Amazonians. Sci. Adv. 6, eaaz1002 (2020).

64. Terrace, H. S. The simultaneous chain: a new approach to serial learning. Trends Cogn. Sci. 9, 202–210 (2005).

65. Brannon, E. M. & Terrace, H. S. Ordering of the Numerosities 1 to 9 by Monkeys. Science 282, 746–749 (1998).

66. Avdagic, E., Jensen, G., Altschul, D. & Terrace, H. S. Rapid cognitive flexibility of rhesus macaques performing psychophysical task-switching. Anim. Cogn. 17, 619–631 (2014).

67. Howard-Spink, E. et al. Nonadjacent dependencies and sequential structure of chimpanzee action during a natural tool-use task. PeerJ 12, e18484 (2024).

68. Cohen, Y. et al. Hidden neural states underlie canary song syntax. Nature 582, 539–544 (2020).

69. Manakov, M. et al. Cognitive Graphs of Latent Structure in Rostral Anterior Cingulate Cortex. 2025.04.18.649507 Preprint at 10.1101/2025.04.18.649507 (2025).

70. El-Gaby, M. et al. A cellular basis for mapping behavioural structure. Nature 1–10 (2024) doi:10.1038/s41586-024-08145-x.

71. Conen, K. E. & Desrochers, T. M. The Neural Basis of Behavioral Sequences in Cortical and Subcortical Circuits. in Oxford Research Encyclopedia of Neuroscience (Oxford University Press, 2022). doi:10.1093/acrefore/9780190264086.013.421.

72. Sternberg, S., Monsell, S., Knoll, R. L. & Wright, C. E. The latency and duration of rapid movement sequences: Comparisons of speech and typewriting. in Information processing in motor control and learning 117–152 (Elsevier, 1978).

73. Wong, K.-F. & Wang, X.-J. A Recurrent Network Mechanism of Time Integration in Perceptual Decisions. J. Neurosci. 26, 1314–1328 (2006).

74. Malassis, R., Dehaene, S. & Fagot, J. Baboons (Papio papio) Process a Context-Free but Not a Context-Sensitive Grammar. Sci. Rep. 10, 7381 (2020).

75. Hong, T. & Stauffer, W. R. Computational complexity drives sustained deliberation. Nat. Neurosci. 26, 850–857 (2023).

76. Schall, J. D. & Thompson, K. G. Neural Selection and Control of Visually Guided Eye Movements. Annu. Rev. Neurosci. 22, 241–259 (1999).

77. Freedman, D. J., Riesenhuber, M., Poggio, T. & Miller, E. K. Categorical Representation of Visual Stimuli in the Primate Prefrontal Cortex. Science 291, 312–316 (2001).

78. Wallis, J. D., Anderson, K. C. & Miller, E. K. Single neurons in prefrontal cortex encode abstract rules. Nature 411, 953–956 (2001).

79. Mansouri, F. A., Freedman, D. J. & Buckley, M. J. Emergence of abstract rules in the primate brain. Nat. Rev. Neurosci. 21, 595–610 (2020).

80. Bernardi, S. et al. The Geometry of Abstraction in the Hippocampus and Prefrontal Cortex. Cell 183, 954–967.e21 (2020).

81. Romo, R., Hernández, A. & Zainos, A. Neuronal Correlates of a Perceptual Decision in Ventral Premotor Cortex. Neuron 41, 165–173 (2004).

82. Okuyama, S., Kuki, T. & Mushiake, H. Recruitment of the premotor cortex during arithmetic operations by the monkey. Sci. Rep. 14, (2024).

83. Wise, S. P. & Murray, E. A. Arbitrary associations between antecedents and actions. Trends Neurosci. 23, 271–276 (2000).

84. Hogeveen, J. et al. What Does the Frontopolar Cortex Contribute to Goal-Directed Cognition and Action? J. Neurosci. 42, 8508–8513 (2022).

85. Xie, Y. et al. Geometry of sequence working memory in macaque prefrontal cortex. Science 375, 632–639 (2022).

86. Petrides, M. Impairments on nonspatial self-ordered and externally ordered working memory tasks after lesions of the mid-dorsal part of the lateral frontal cortex in the monkey. J. Neurosci. 15, 359– 375 (1995).

87. Fujii, N. Representation of Action Sequence Boundaries by Macaque Prefrontal Cortical Neurons.Science 301, 1246–1249 (2003).

88. Ohbayashi, M., Picard, N. & Strick, P. L. Inactivation of the Dorsal Premotor Area Disrupts Internally Generated, But Not Visually Guided, Sequential Movements. J. Neurosci. 36, 1971–1976 (2016).

89. Russo, A. A. et al. Neural Trajectories in the Supplementary Motor Area and Motor Cortex Exhibit Distinct Geometries, Compatible with Different Classes of Computation. Neuron 107, 745–758.e6 (2020).

90. Shima, K. & Tanji, J. Neuronal Activity in the Supplementary and Presupplementary Motor Areas for Temporal Organization of Multiple Movements. J. Neurophysiol. 84, 2148–2160 (2000).

91. Lu, X. & Ashe, J. Anticipatory Activity in Primary Motor Cortex Codes Memorized Movement Sequences. Neuron 45, 967–973 (2005).

92. Matsuzaka, Y., Picard, N. & Strick, P. L. Skill Representation in the Primary Motor Cortex After Long-Term Practice. J. Neurophysiol. 97, 1819–1832 (2007).

93. Fuster, J. M. Upper processing stages of the perception–action cycle. Trends Cogn. Sci. 8, 143– 145 (2004).

94. Fodor, J. A. & Pylyshyn, Z. W. Connectionism and cognitive architecture: A critical analysis. Cognition 28, 3–71 (1988).

95. Barbas, H. & Pandya, D. N. Architecture and frontal cortical connections of the premotor cortex (area 6) in the rhesus monkey. J. Comp. Neurol. 256, 211–228 (1987).

96. Rizzolatti, G., Luppino, G. & di Parma, U. The Cortical Motor System Review. (2001).

97. Mahowald, K. et al. Dissociating language and thought in large language models. Trends Cogn. Sci. 28, 517–540 (2024).

98. Lee, A., Viégas, F. & Wattenberg, M. Tensor Product Representation Probes Reveal Shared Structure Across Linear Directions. Preprint at 10.48550/arXiv.2605.09967 (2026).

99. Orlov, T., Yakovlev, V., Hochstein, S. & Zohary, E. Macaque monkeys categorize images by their ordinal number. Nature 404, 77–80 (2000).

100. Penn, D. C., Holyoak, K. J. & Povinelli, D. J. Darwin’s mistake: Explaining the discontinuity between human and nonhuman minds. Behav. Brain Sci. 31, 109–130 (2008).

101. Sohn, J.-W. & Lee, D. Order-Dependent Modulation of Directional Signals in the Supplementary and Presupplementary Motor Areas. J. Neurosci. 27, 13655–13666 (2007).

102. Isoda, M. & Tanji, J. Cellular Activity in the Supplementary Eye Field During Sequential Performance of Multiple Saccades. J. Neurophysiol. 88, 3541–3545 (2002).

103. Procyk, E., Tanaka, Y. L. & Joseph, J. P. Anterior cingulate activity during routine and non-routine sequential behaviors in macaques. Nat. Neurosci. 3, 502–508 (2000).

104. Kermadi, I. & Joseph, J. P. Activity in the caudate nucleus of monkey during spatial sequencing. J. Neurophysiol. 74, 911–933 (1995).

105. Mendoza, G., Méndez, J. C., Pérez, O., Prado, L. & Merchant, H. Neural basis for categorical boundaries in the primate pre-SMA during relative categorization of time intervals. Nat. Commun. 9, 1098 (2018).

106. Altschul, D., Jensen, G. & Terrace, H. Perceptual category learning of photographic and painterly stimuli in rhesus macaques (Macaca mulatta) and humans. PLOS ONE 12, e0185576 (2017).

107. Nieder, A. & Dehaene, S. Representation of Number in the Brain. Annu. Rev. Neurosci. 32, 185– 208 (2009).

108. Makino, Y., Suemitsu, K. & Hirashima, M. Action Progress Provides an Abstract Coordinate for Motor Memory. 2026.02.09.704807 Preprint at 10.64898/2026.02.09.704807 (2026).

109. Marcus, G. F. Rethinking Eliminative Connectionism. Cognit. Psychol. 37, 243–282 (1998).

110. Remington, E. D., Narain, D., Hosseini, E. A. & Jazayeri, M. Flexible Sensorimotor Computations through Rapid Reconfiguration of Cortical Dynamics. Neuron 98, 1005–1019.e5 (2018).

111. Akkal, D., Bioulac, B., Audin, J. & Burbaud, P. Comparison of neuronal activity in the rostral supplementary and cingulate motor areas during a task with cognitive and motor demands. Eur. J. Neurosci. 15, 887–904 (2002).

112. Clower, W. T. & Alexander, G. E. Movement Sequence-Related Activity Reflecting Numerical Order of Components in Supplementary and Presupplementary Motor Areas. J. Neurophysiol. 80, 1562–1566 (1998).

113. Mushiake, H., Inase, M. & Tanji, J. Selective coding of motor sequence in the supplementary motor area of the monkey cerebral cortex. Exp. Brain Res. 82, (1990).

114. Nakajima, T., Hosaka, R., Mushiake, H. & Tanji, J. Covert Representation of Second-Next Movement in the Pre-Supplementary Motor Area of Monkeys. J. Neurophysiol. 101, 1883–1889 (2009).

115. Nachev, P., Kennard, C. & Husain, M. Functional role of the supplementary and pre-supplementary motor areas. Nat. Rev. Neurosci. 9, 856–869 (2008).

116. Mita, A., Mushiake, H., Shima, K., Matsuzaka, Y. & Tanji, J. Interval time coding by neurons in the presupplementary and supplementary motor areas. Nat. Neurosci. 12, 502–507 (2009).

117. Merchant, H. et al. Different time encoding strategies within the medial premotor areas of the primate. Preprint at 10.1101/2023.01.28.526038 (2023).

118. Shima, K., Mushiake, H., Saito, N. & Tanji, J. Role for cells in the presupplementary motor area in updating motor plans. Proc. Natl. Acad. Sci. 93, 8694–8698 (1996).

119. Isoda, M. & Hikosaka, O. Switching from automatic to controlled action by monkey medial frontal cortex. Nat. Neurosci. 10, 240–248 (2007).

120. Nakamura, K., Sakai, K. & Hikosaka, O. Neuronal Activity in Medial Frontal Cortex During Learning of Sequential Procedures. J. Neurophysiol. 80, 2671–2687 (1998).

121. Shima, K. & Tanji, J. Both Supplementary and Presupplementary Motor Areas Are Crucial for the Temporal Organization of Multiple Movements. J. Neurophysiol. 80, 3247–3260 (1998).

122. Halsband, U., Matsuzaka, Y. & Tanji, J. Neuronal activity in the primate supplementary, pre-supplementary and premotor cortex during externally and internally instructed sequential movements. Neurosci. Res. 20, 149–155 (1994).

123. Cona, G. & Semenza, C. Supplementary motor area as key structure for domain-general sequence processing: A unified account. Neurosci. Biobehav. Rev. 72, 28–42 (2017).

124. Levy, D. et al. Role for Left Dorsomedial Prefrontal Cortex in Self-Generated, but not Externally Cued, Language Production. Neurobiol. Lang. 6, nol_a_00166 (2025).

125. Bengtsson, S. L., Csíkszentmihályi, M. & Ullén, F. Cortical Regions Involved in the Generation of Musical Structures during Improvisation in Pianists. J. Cogn. Neurosci. 19, 830–842 (2007).

126. Chen, Q., Beaty, R. E. & Qiu, J. Mapping the artistic brain: Common and distinct neural activations associated with musical, drawing, and literary creativity. Hum. Brain Mapp. 41, 3403–3419 (2020).

127. Planton, S., Jucla, M., Roux, F.-E. & Démonet, J.-F. The “handwriting brain”: A meta-analysis of neuroimaging studies of motor versus orthographic processes. Cortex 49, 2772–2787 (2013).

128. Leek, E. C., Yuen, K. S. L. & Johnston, S. J. Domain General Sequence Operations Contribute to Pre-SMA Involvement in Visuo-spatial Processing. Front. Hum. Neurosci. 10, (2016).

129. Fehr, T., Code, C. & Herrmann, M. Common brain regions underlying different arithmetic operations as revealed by conjunct fMRI–BOLD activation. Brain Res. 1172, 93–102 (2007).

130. Hanakawa, T. et al. The Role of Rostral Brodmann Area 6 in Mental-operation Tasks: an Integrative Neuroimaging Approach. Cereb. Cortex 12, 1157–1170 (2002).

131. Kean, H. et al. A human brain network specialized for abstract formal reasoning. 2025.10.21.683445 Preprint at 10.1101/2025.10.21.683445 (2025).

132. Ninokura, Y., Mushiake, H. & Tanji, J. Representation of the Temporal Order of Visual Objects in the Primate Lateral Prefrontal Cortex. J. Neurophysiol. 89, 2868–2873 (2003).

133. Tanji, J. & Hoshi, E. Role of the Lateral Prefrontal Cortex in Executive Behavioral Control. Physiol. Rev. 88, 37–57 (2008).

134. Shallice, T. & Burgess, P. W. DEFICITS IN STRATEGY APPLICATION FOLLOWING FRONTAL LOBE DAMAGE IN MAN. Brain 114, 727–741 (1991).

135. Histed, M. H. & Miller, E. K. Microstimulation of Frontal Cortex Can Reorder a Remembered Spatial Sequence. PLOS Biol. 4, e134 (2006).

136. Tian, L. Y. & Brainard, M. S. Discrete Circuits Support Generalized versus Context-Specific Vocal Learning in the Songbird. Neuron 96, 1168–1177.e5 (2017).

137. Ohbayashi, M. The Roles of the Cortical Motor Areas in Sequential Movements. Front. Behav. Neurosci. 15, (2021).

138. Mills, T., et al. Spatiotemporal Program Learning in Human Adults, Children, and Monkeys. Preprint at 10.31234/osf.io/2n85j_v2 (2025).

139. Zylberberg, A., Dehaene, S., Roelfsema, P. R. & Sigman, M. The human Turing machine: a neural framework for mental programs. Trends Cogn. Sci. S136466131100088X (2011) doi:10.1016/j.tics.2011.05.007.

140. Liu, Y., Tian, L. Y., Tenenbaum, J. B., Freiwald, W. A. & Wang, X.-J. A neural circuit algorithm for action grammar. 2026.10.06.757194 Preprint at 10.64898/2026.10.06.757194 (2026).

141. McClelland, J. L. et al. Letting structure emerge: connectionist and dynamical systems approaches to cognition. Trends Cogn. Sci. 14, 348–356 (2010).

142. Gelder, T. van. The dynamical hypothesis in cognitive science. Behav. Brain Sci. 21, 615–628 (1998).

143. Mollazadeh, M. et al. Spatiotemporal Variation of Multiple Neurophysiological Signals in the Primary Motor Cortex during Dexterous Reach-to-Grasp Movements. J. Neurosci. 31, 15531– 15543 (2011).

144. Saleem, K. S. & Logothetis, N. K. A Combined MRI and Histology Atlas of the Rhesus Monkey Brain in Stereotaxic Coordinates. (Academic Press, 2012).

145. He, S., Dum, R. & Strick, P. Topographic organization of corticospinal projections from the frontal lobe: motor areas on the medial surface of the hemisphere. J. Neurosci. 15, 3284–3306 (1995).

146. Hwang, J., Mitz, A. R. & Murray, E. A. NIMH MonkeyLogic: Behavioral control and data acquisition in MATLAB. J. Neurosci. Methods 323, 13–21 (2019).

147. Pachitariu, M., Sridhar, S., Pennington, J. & Stringer, C. Spike sorting with Kilosort4. Nat. Methods 21, 914–921 (2024).

148. Testard, C. et al. Neural signatures of natural behaviour in socializing macaques. Nature 628, 381–390 (2024).

149. Mante, V., Sussillo, D., Shenoy, K. V. & Newsome, W. T. Context-dependent computation by recurrent dynamics in prefrontal cortex. Nature 503, 78–84 (2013).

150. Liu, Y., Brincat, S. L., Miller, E. K. & Hasselmo, M. E. A Geometric Characterization of Population Coding in the Prefrontal Cortex and Hippocampus during a Paired-Associate Learning Task. J. Cogn. Neurosci. 32, 1455–1465 (2020).

151. Willett, F. R. et al. Hand Knob Area of Premotor Cortex Represents the Whole Body in a Compositional Way. Cell 181, 396–409.e26 (2020).

